# Phylogenetic analysis of enteroviruses from non-human primates reveals two new species within the genus *Enterovirus* and inter-species recombination

**DOI:** 10.64898/2026.01.19.700265

**Authors:** Corentin Aubé, Paulina Cruz de Casas, Matthieu Prot, Artem Baidaliuk, Marie-Claire Endegue Zanga, Etienne Simon-Lorière, Nolwenn Jouvenet, Serge A. Sadeuh-Mba, Maël Bessaud

## Abstract

To date, 15 species have been described within the genus *Enterovirus.* Previous studies suggested the existence of another species comprising strains isolated from the stool specimens of non-human primates (NHPs) in Central Africa. Moreover, numerous full-length or partial genomic sequences of NHP enteroviruses (EVs) can be found in GenBank without being properly classified. To our knowledge, no comprehensive synthesis of NHP EV data exists, leaving genetic relationships between strains across independent studies unclear. To address these gaps, we sequenced the complete genome of four NHP EVs from our stool collection and conducted an extensive search of NHP EV sequences in GenBank to perform a comprehensive phylogenetic analysis. Our analyses revealed two new species tentatively named *Enterovirus mbel* and *Enterovirus noa*, which contain at least 6 and 2 virus types, respectively. We also identified new virus types within the known species EV-J.

Phylogenetic analyses strongly suggest interspecies recombination events between NHP EVs in the non-structural region of the genome, challenging the long-held view that recombination is confined within narrowly defined subsets of EVs belonging to the same species. We also performed the first comprehensive comparative analysis of full length human and NHP EV genomes, focusing on GC content, dinucleotide frequencies and codon-usage bias. GC content emerged as the most robust host-associated marker: all NHP-associated virus types within species *E. alphacoxsackie* and *E. betacoxsackie* displayed GC % below 47 %, whereas human-derived virus types exhibited GC % above 47 %. Dinucleotide frequency, Effective Number of Codons (ENC) and Relative Synonymous Codon Usage highlighted distinct codon-bias clusters that mirror the phylogenetic relationships between EVs but only partially correlate with their respective hosts of origin.

This work enhances our understanding of EVs circulating in NHPs and paves the way for future research aiming at understanding the mechanisms underlying host-adaptation among EVs.

## INTRODUCTION

Enteroviruses (EVs) are small, non-enveloped viruses with an icosahedral capsid. Their genome is a single-stranded and positive-sense RNA molecule of approximately 7,200-8,500 nucleotides (nt) in length. It contains two untranslated regions (5’UTR and 3’UTR) flanking a large open reading frame (ORF) that encodes a polyprotein, which is subsequently cleaved into three precursors (P1, P2 and P3). P1 gives rise to the four capsid proteins (VP1 to VP4) while P2 and P3 are cleaved into different proteins involved in viral replication. A short additional upstream ORF (uORF) encoding a single protein has been described in the genome of some EVs (Lulla et al., 2019).

The classification of EVs has undergone significant revisions since their discovery. Initially, EVs were classified into different groups based on their phenotypic properties (e.g. tropism and pathogenicity in mice) and physicochemical characteristics (density of the virus particles, sensibility to acid pHs) (Bessaud et al., 2018). Within each group, virus isolates were grouped into serotypes, based on *in vitro* cross-neutralization assays. In the 2000s, genetic criteria replaced phenotypic and serological criteria as the basis for classification (M. S. Oberste et al., 1999). Enteroviruses are now classified into species and virus types, depending on their phylogenetic relationships. According to the International Committee for Taxonomy of Viruses guidelines (ICTV), EVs belong to the same species if they share at least 70% amino acid (aa) identity in the complete ORF and in the 2C+3CD genes, and at least 60% aa identity in the capsid (P1 region). The *Enterovirus* genus currently comprises 15 species: *Enterovirus alphacoxsackie* (EV-A), *Enterovirus betacoxsackie* (EV-B), *Enterovirus coxsackiepol* (EV-C), *Enterovirus deconjuncti* (EV-D) *Enterovirus eibovi* (EV-E), *Enterovirus fitauri* (EV-F) *Enterovirus geswini* (EV-G), *Enterovirus hesimi* (EV-H), *Enterovirus idromi* (EV-I), *Enterovirus jesimi* (EV-J), *Enterovirus krodeni* (EV-K), *Enterovirus lesimi* (EV-L), *Enterovirus alpharhino* (RV-A), *Enterovirus betarhino* (RV-B) and *Enterovirus cerhino* (RV-C), each of which comprise multiple virus types based on the VP1-encoding sequence. Virus types within a species are defined by >75% nt identity and >85%-88% aa identity in this region (Brown et al., 2009; M. Steven Oberste et al., 1999).

EVs infect a wide range of mammals, including humans, non-human primates (NHPs), small ruminants, pigs, cattle, camels and rodents. While each EV appears adapted to a limited set of host species that constitutes its natural reservoirs, host barriers can be overcome, as demonstrated by numerous documented cross-species transmissions between humans and animals, bidirectionally, or among different animal species (Doté et al., 2024; Fieldhouse et al., 2018; Modiyinji et al., 2025). Notably, EVs do not exhibit clear host-associated phylogenetic clustering, as some species include virus types circulating in different hosts. For instance, some EV-As, EV-Bs and EV-Ds circulate in humans while others circulate in NHPs. Similarly, some EV-Gs circulate in pigs while others circulate in small ruminants.

The first identified NHP EVs were detected in simian cell cultures and laboratory animals in the mid-20^th^ century (Oberste et al., 2002). Since then, new EV types have been discovered in captive, wild, healthy and sick NHPs. Over the past 20 years, next-generation sequencing has increased the number of publicly available NHP EV sequences, however, data remain sparse. Furthermore, many sequences span diverse sub-genomic regions and remain to be properly classified. Moreover, no comprehensive synthesis of all available NHP EV molecular data exists, leaving the genetic relationships between NHP EVs across independent studies fragmented. Additionally, sequences of NHP EVs sampled in Central Africa have suggested the existence of at least one additional EV species (Sadeuh-Mba et al., 2014), but it has never been formally described. To address these gaps, we conducted an extensive search of NHP EV sequences in GenBank and performed a comprehensive phylogenic analysis. To make this analysis as complete as possible, we also sequenced the complete genome of four NHP EVs from our NHP stool collection and samples from our previous work (Sadeuh-Mba et al., 2014). Furthermore, we compared genetic characteristics, such as GC percentage (GC%), dinucleotide ratios or codon usage (effective number of codons, ENC and relative synonymous codon usage, RSCU) between NHP and human EVs. Our analysis identified two previously uncharacterized EV species and revealed that specific genomic signatures align with phylogeny and may be associated with host-adaptation.

## METHODS

### Amplicon-based sequencing

EV-B114 strain Z057 was grown in RD cells and RNA was extracted from 250 µL of clarified cell culture supernatant using the High Pure Viral RNA kit (Roche Diagnostics, Meylan, France) following the manufacturer’s instructions. Genome amplification was performed by RT-PCR with primers targeting previously described conserved regions of the EV genome (Joffret et al., 2018). Sequencing was performed by Illumina technology and de novo assembly conducted with CLC Genomics Workbench as previously described (Bessaud et al., 2016). As EV-J121 strain CHE20 could not be propagated in cell culture (Sadeuh-Mba et al., 2014), RNA was extracted directly from 140 µL of clarified 20% (Weight/Volume) stool suspension using the QIAamp Viral RNA Mini Kit (Qiagen, France). Genome amplification was performed by RT-PCR with primers targeting the sub-genomic regions previously sequenced (Sadeuh-Mba et al., 2014), as well as a primer targeting the 3’ terminal poly A sequence. Sequencing was performed by the Sanger technique.

### Metatranscriptomic next-generation sequencing

Two strains, RCMH3 and RCMH6, that could not be grown in cell culture were sequenced using a protocol previously described (Gámbaro et al., 2021). After extraction from 140 μl of fecal suspension using the QIAamp Viral RNA Mini Kit (Qiagen) followed by Turbo DNAse treatment (Ambion), RNA was purifed with Agencourt RNAClean XP beads. Host rRNA were depleted using custom probes and RNAse H treatment (Matranga et al., 2014). After purification using AMPure RNA clean beads (Beckman Coulter Genomics) and elution in 10 μl of RNase-free water, RNA was retro transcribed using random primers and SuperScript IV (Invitrogen). Second-strand DNA was synthesized using *E. coli* DNA ligase, RNAse H and DNA polymerase (New England Biolabs) and purified using Agencourt AMPure XP beads (Beckman Coulter). Libraries were then prepared using the Nextera XT DNA Library Prep Kit (Illumina), quality controlled using a Bioanalyzer high sensitivity DNA analysis kit (Agilent) and sequenced on an Illumina NextSeq500 (2 × 75 cycles).

Adapters and low-quality reads were removed using Trimmomatic v0.39 (Bolger et al., 2014). Trimmed reads were assembled using megahit v1.2.9 (Li et al., 2016) with default parameters. The contigs were queried against the NCBI non-redundant protein database using DIAMOND v2.0.4 (Buchfink et al., 2015), leading to the identification of novel EV sequences. The EV genomes were then reconstructed by iterative mapping of the trimmed reads, using clc-assembly-cell v5.1.0. We used Samtools v1.10 (Li et al., 2009) to generate alignment statistics, and ivar v1.0 (Grubaugh et al., 2019) to generate consensus sequences, using a minimum of 3× read depth coverage. The mapping data was visually checked to confirm the accuracy of the obtained genomes using Geneious Prime 2023 (www.geneious.com).

### Phylogenetic analysis

Multiple sequence alignments were performed with CLC Main Workbench 24.0.2 software (CLC bio, Aarhus, Denmark). Phylogenetic trees were inferred by the neighbour-joining (NJ) or maximum likelihood (ML) methods implemented in MEGA X (Kumar et al., 2018). Distance-based phylogenetic trees were reconstructed by the NJ method with the Tajima-Nei model for computing evolutionary distances (Tajima and Nei, 1984). The ML phylogenetic trees were reconstructed by the Tajima-Nei model and Nearest-Neighbor-Interchange method for ML heuristic method. The reliability of tree topologies was estimated by bootstrap analysis with 1,000 pseudoreplicates for NJ trees and 500 pseudoreplicates for ML trees due to computational constraints. Tree representations were generated using MEGA X (version 10.2.6) or FigTree (version 1.4.4) (https://tree.bio.ed.ac.uk/software/figtree/). Links to Nextstrain interactive representations of trees from Figures 2 and 6, Supplementary Figure 1, and tanglegrams are available on GitHub: https://github.com/CorentinAube/NHP_EV_genomic_analysis.

### Genomic comparison

Prototype genomes of human and NHP EVs were retrieved from the Picornaviridae database (http://www.picornaviridae.com), while more recent sequences were retrieved from the BV-BRC database (Olson et al., 2023). Other partial NHP EV sequences were retrieved from the scientific literature indexed in PubMed. Protein sequence translation, MSA, nucleotide and amino acid identity matrices were calculated with CLC Main Workbench. The GC content, dinucleotide ratios, ENC and RSCU values were calculated using custom Python (version 3.12.12) scripts available on GitHub with also the list of all accession numbers of sequences used (https://github.com/CorentinAube/NHP_EV_genomic_analysis). The formulas for dinucleotide ratio, ENC and RSCU calculations were obtained from Zeng and colleagues (Zeng et al., 2022). The determination of GC3 bias was based on the method published by Butt and colleagues (Butt et al., 2016).

### Data availability

Nucleotide sequences generated in this study were deposited on the GenBank database (accession numbers PX682287-PX682290). An isolate of the EV-B114 strain Z057 has been shared via the EVAg repository (Romette et al., 2018).

## RESULTS

### Whole-genome sequencing of four NHP EVs from our stool collection

Four EVs that had only been previously partially sequenced (Sadeuh-Mba et al., 2014), were subjected to whole genome sequencing. Contigs covering nearly the entire viral genome were obtained for three viruses: Z057, CHE20, and RCMH3 (Figure 1A). For RCMH6, approximately 150 nt could not successfully be sequenced at the 5’ extremity. All 4 viruses featured the canonical EV genomic organization, with a large ORF flanked by two UTRs. Comparing these sequences with annotated reference EV genomes, canonical polyprotein cleavage sites were identified, suggesting that the polyprotein processing in these viruses is similar to that of other EVs (Supplementary Table 1). Additionally, the genomes of Z057, RCMH3, and RCMH6 contained a putative uORF. This putative uORF overlaps with the main ORF with a frameshift, and its putative start codon is located 133 or 136 nt upstream of the main ORF (Figure 1B). In contrast, the EV-J121 CHE20 genome lacked a uORF, as previously reported for other members of the EV-J species (Lulla et al., 2019). All four genomes also contained a cis-replication element (CRE) within the 2C gene, matching the consensus sequence 5’-GXXXAAAXXXXXXA-3’ (Figure 1C) (Cordey et al., 2008).

**Figure 1.**
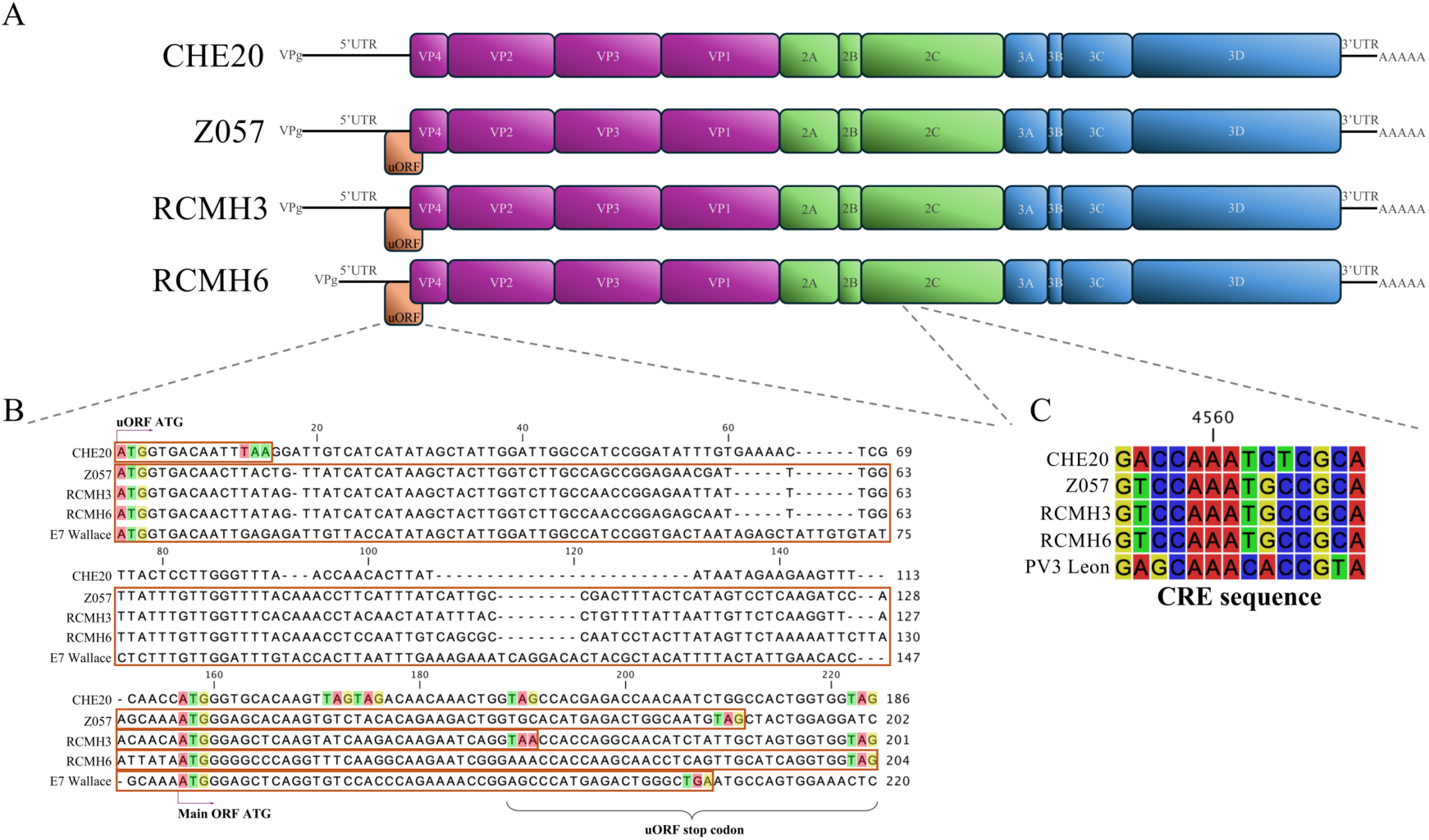
Genome organization of CHE20, Z057, RCMH6, and RCMH3 strains. **(A)** Schematic representation of the genomic organization of the four newly sequenced genomes. The uORF is shown in brown (Lulla et al., 2019), the P1 region in purple, the P2 region in green, and the P3 region in blue. **(B)** Alignment of nucleotide sequences of the potential uORF compared to the E7 Wallace strain (highlighted by brown boxes). The start codons of the putative uORF and the main ORF, as well as, the stop codons of the uORF are highlighted. **(C)** Alignment of the CRE sequence compared to the PV3 Leon strain (Cordey et al., 2008). The sequence alignment was performed using the CLC Main Workbench v24 software.

### Comprehensive phylogeny of non-human primate enteroviruses

The search for NHP EV sequences in GenBank led to the identification of 336 sequences, of which 24 were complete genomes, 91 were VP1 sequences, 103 were VP4/2 sequences, 19 were VP2 sequences, 26 were 3D sequences and 29 were sequences covering various regions of the genome, referred to as partial (Table 1).

**Table 1.**
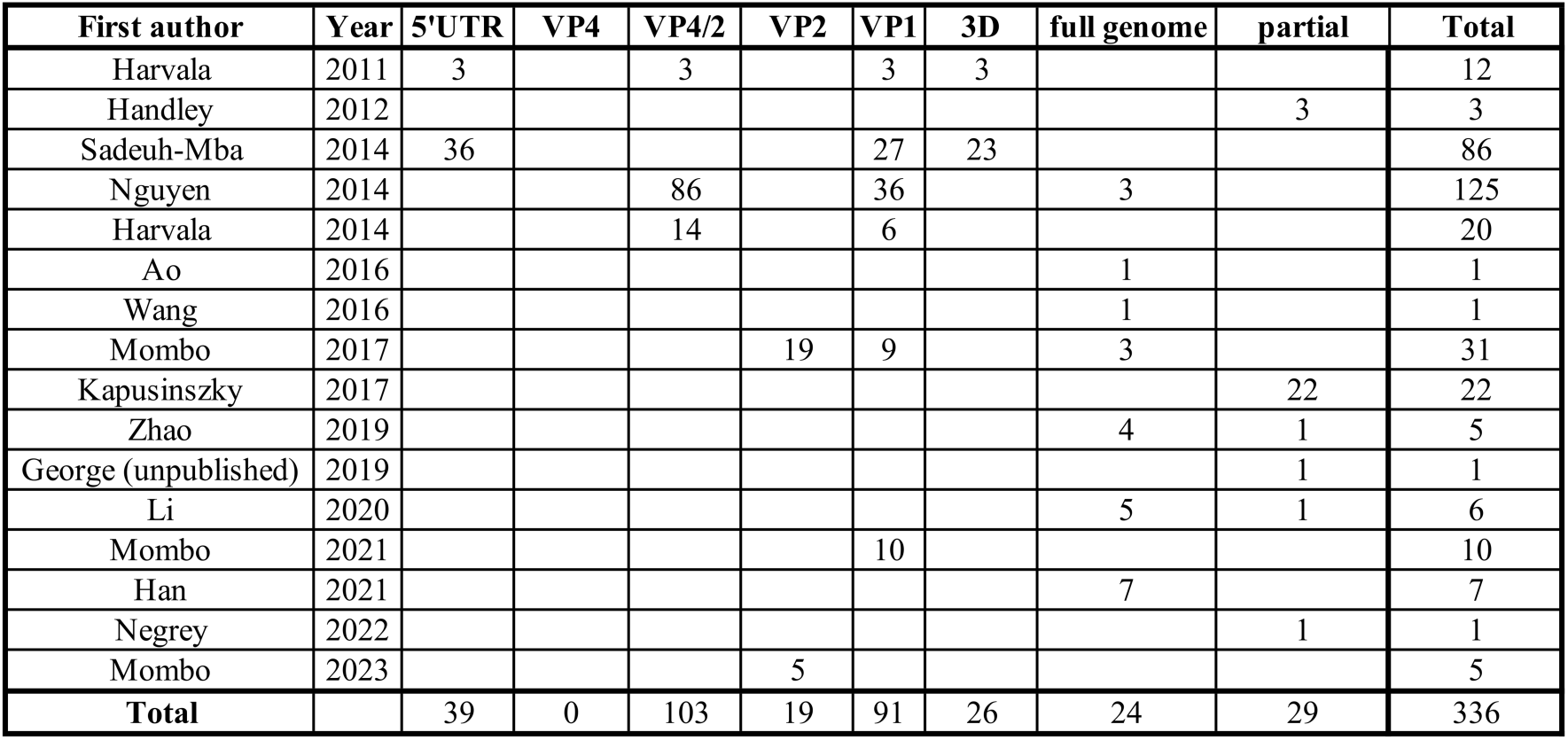
Summary of included studies and number of available enterovirus sequences. For each study, the first author, year of publication, and the number of sequences available are listed. Data were obtained from: (Ao et al., 2016; Han et al., 2021; Handley et al., 2012; Harvala et al., 2011, 2014; Kapusinszky et al., 2017; Li et al., 2020; Mombo et al., 2017, 2021, 2023; Negrey et al., 2022; Nguyen et al., 2014; Sadeuh-Mba et al., 2014; Wang et al., 2016; Zhao et al., 2019)

A phylogenetic tree based on the complete VP1 gene was generated using all available NHP EV sequences spanning this region along with representatives of all known human derived EV types (Figure 2 and Supplementary Figure 1). This VP1-based tree clearly delineates the EV species. Non-human primate derived EVs were found in species EV-A, -B, -D, -H, -J and - L. The data for EV-L are very sparse with only one known representative. Species EV-H includes only 6 sequences, of which 4 are epidemiologically related to each other. To determine whether these sequences belong to the same virus type, the 75% nt and 85% aa identity rule (Brown et al., 2009) was applied to their VP1 sequences. As previously described (Nguyen et al., 2014), these sequences belong to a single virus type (EV-H1), sharing 84.09%-100% nt identity and 96.17%-100% aa identity (Supplementary Figure 2). In contrast to the EV-H and EV-L species, distinct EV-J virus types have been previously described. The capsid-based phylogram accurately delineated the EV-J103, EV-J108 and EV-J122 virus types and confirmed the EV-J121 type, of which CHE20 is the first known representative that was previously characterized based on partial sequence (Sadeuh-Mba et al., 2014) (Figure 3A). In the VP4/VP2 region, three strains (CPML8127, CPOV3340 and CPOV3342) were closely related to CHE20 (Figure 3C), thus suggesting that they could also be assigned to the EV-J121 virus type. Identity matrices were calculated to assign several untyped EV-J sequences (Figure 3A and 3B, Supplementary Figure 3). NOLA1 and NOLA3 share over 75.41% and 85.31% at nt and aa level, respectively, with an EV-J103 strain and were thus assigned to this virus type. Similarly, the sequence comparison of the partial VP1 sequence of cg5275 with EV-J prototype strains supported its assignment to the EV-J103 type (>77.36% and >85.43 nt and aa identity, respectively, with other EV-J103 viruses, Supplementary Figure 4). As previously reported (Wang et al., 2016) the Sev-nJ strain represent a novel virus type, sharing less than 74.48% and 84.27% with other EV-J sequences (Supplementary Figure 3). The CHN-BJ-LXL1 sequence, published 5 years later than the Sev-nJ strain, belonged to the same EV-J type, as it shares 82.09% and 92.63% nt and aa identity with Sev-nJ, respectively (Supplementary Figure 3). We propose to assign these two strains to a novel virus type, EV-J123, following current taxonomic nomenclature. Additionally, several partial VP1 and VP4/2 sequences remained to be assigned (Harvala et al., 2014). Based on partial VP1 sequences, the CPOV3336 strain is related to EV-J121 strain CHE20 (Figure 3B), but their nt and aa identities (66.04% and 75.81%, Supplementary Figure 4) indicating that CPOV3336 belongs to a putative distinct virus type (tentatively named EV-J124). A set of short VP4/VP2 sequences (CPOV3337, DJ4139, CPGR7921, CPLM8102 and CPLM8125) were very close that of CPOV3336 (Figure 3C), suggesting that they belong to the newly proposed EV-J124 type.

**Figure 2.**
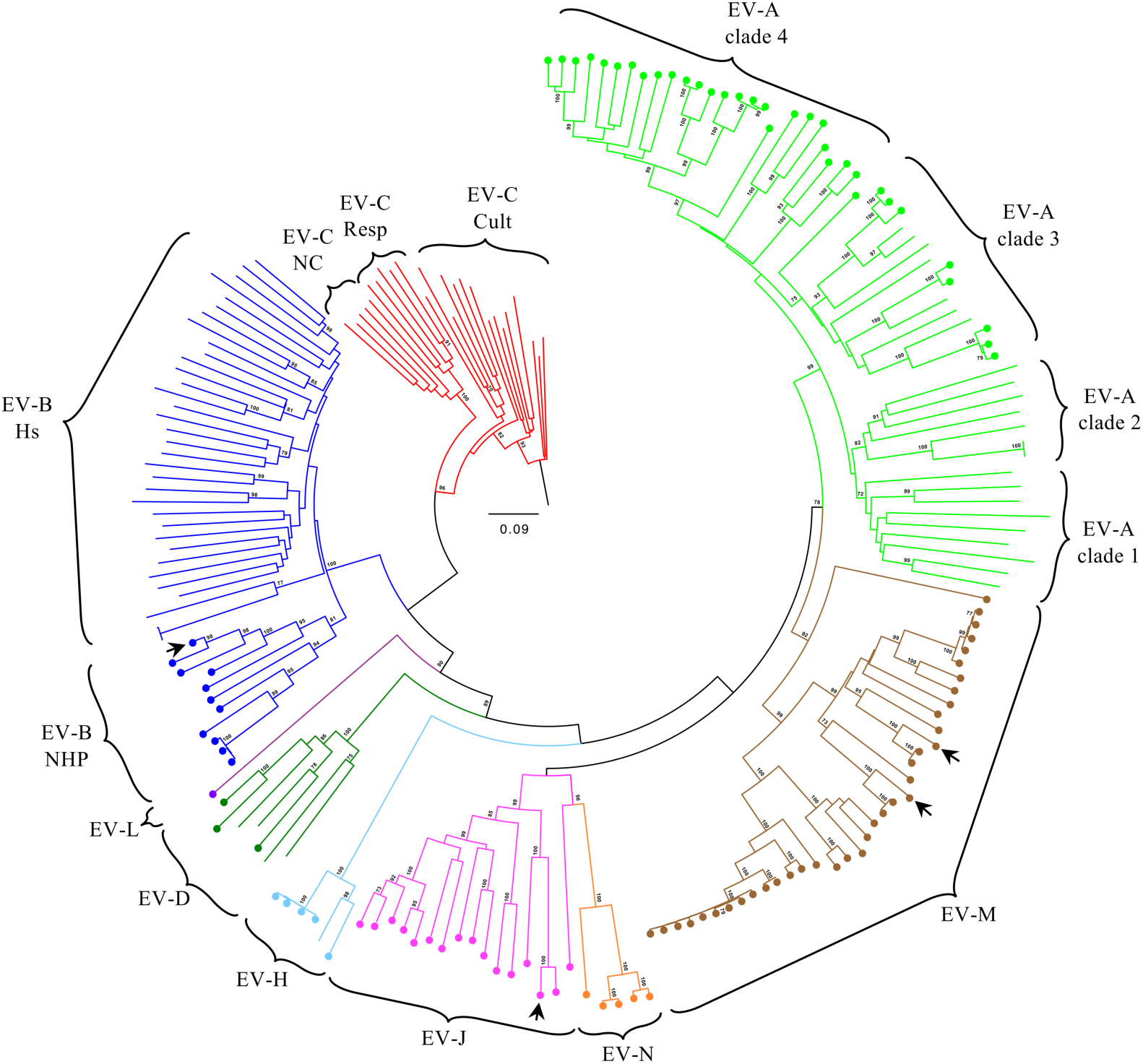
Phylogenetic analysis of complete VP1 sequences from non-human primate and representative human enteroviruses. The Neighbor-Joining tree was reconstructed from a full-length VP1 nucleotide sequence alignment. Branches are coloured according to EV species: EV-A (light green), EV-B (dark blue), EV-C (red), EV-D (dark green), EV-H (light blue), EV-J (pink), EV-L (purple), EV-M (brown) and EV-N (orange). Circles indicate NHP derived EV strains. The arrows indicate new genomes sequenced in our study (Figure 1). The tree is rooted with the PV1 Mahoney strain (V01148). Bootstrap values greater than 70% (based on 1,000 pseudoreplicates) are shown at the corresponding nodes. The sequence alignment was performed using the CLC Main Workbench v24, the phylogenetic tree was constructed with MEGA X and the visualization was generated with FigTree. The tree is drawn to scale, with branch lengths measured in the number of substitutions per site.

**Figure 3.**
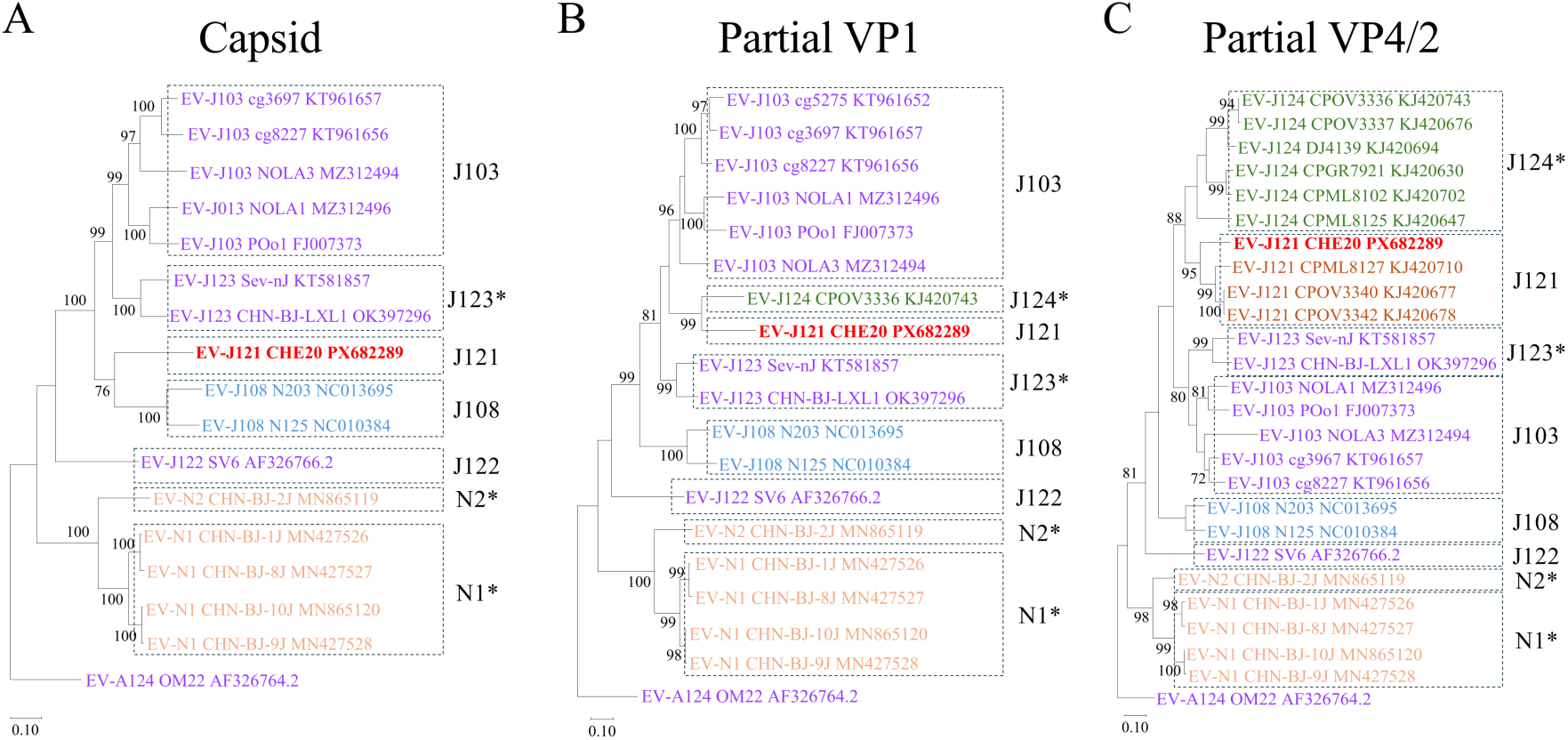
Phylogenetic analysis of enterovirus J species. The maximum-likelihood trees were constructed with all available full capsid (length of alignment 2571 nt) **(A)**, partial VP1 (length of alignment 754 nt) **(B)** and partial VP4/2 (length of alignment 429 nt) **(C)** sequences. The trees were rooted with the EV-A124 OM22 strain. The symbol * indicates proposed new type classifications. The colours of the strain names indicate their host origins: *Mandrillus* (green), *Chlorocebus* (orange), *Cercocebus* (brown), *Papio* (blue), *Macaca* (purple) and *Pan* (red). Bootstrap values greater than 70% (based on 500 pseudoreplicates) are shown at the corresponding nodes. The sequence alignments were performed using the CLC Main Workbench v24, the phylogenetic trees were constructed with MEGA X and the visualization was generated with FigTree. The trees are drawn to scale, with branch lengths measured in the number of substitutions per site.

Some viruses previously identified as EV-Js (CHN-BJ-2J, -1J, -8J, -10J and -9J) form a specific group (Figure 3). To determine whether these viruses defined a separated additional new species, we applied the ICTV criteria. The CHN-BJ-2J, 1J, 8J, 10J and 9J sequences share 69.41% to 72.74% identity in the ORF and 80.72% to 84.50% in the 2C+3CD genes with other EV-J types (Supplementary Figure 5A and B). However, they share only between 55.36% to 59.25% aa identity in the capsid gene (Supplementary Figure 5C), as previously reported (Li et al., 2020). These results support the existence of a separated new species, which we tentatively named *Enterovirus nao* (EV-N), after the Chinese word náo (猱) meaning “monkey” (Couvreur, 1896; Loewe, 1993). As previously suggested (Li et al., 2020), CHN-BJ-1J, 8J, 9J, 10J and CHN-BJ-1-2J form two distinct EV types within this species, as they share less than 72.94% nt and 80.99% aa identity (Supplementary Figure 3). We propose naming these types EV-N1 (for the CHN-BJ-1J, 8J, 9J, 10J strains) and EV-N2 (for the CHN-BJ-2J strain).

Three EV species, EV-A, -B and -D, include some EVs that originated from humans while others were recovered from NHPs. Within the EV-B species, NHP EVs segregate from human derived EVs and define a specific branch (Figure 2). The unique exception to the host association of EV-B types is the case of GAB98 strain, isolated from wild chimpanzees, which is an EV-B107 known to circulate in humans in Asia (Maan et al., 2019; Yamashita et al., 2010). The isolation of strain GAB98 in a chimpanzee was explained as a sporadic human-to-NHP transmission, potentially facilitated by the adjacent or overlapping humans and chimpanzee habitats (Mombo et al., 2017). As previously reported, 4 distinct virus types exist in the NHP specific branch of the EV-B species. Strain Z057, of which a complete genome sequence was obtained in this work (Figure 1), belongs to the EV-B114 virus type (Figure 4), that was first isolated in 1963 from NHP derived cell cultures (Malherbe and Harwin, 1963). Since its discovery, only two EV-B114 EVs have been reported: CPOV3344 (Nguyen et al., 2014) and Z057 (Sadeuh-Mba et al., 2014). They share 82.09%-84.58% nt and 92.88%-93.95% aa identities between each other in the VP1 sequence; thus confirming that they belong to the same virus type (Figure 4). In contrast to EV-B species, the phylogenetic boundaries between NHP and human EVs are less distinct in species EV-A and EV-D (Figure 2). As previously reported, members of the species EV-A fall into four main phylogroups. Clades 1 and 2 consist of EV types that circulate in humans worldwide (Clade 1: EV-A120, EV-A71, CVA7, CVA14 and CVA16; Clade 2: EV-A114, CVA2, CVA3, CVA4, CVA5, CVA6, CVA8, CVA10 and CVA12). Clade 3 includes EVs detected both in human and NHPs (Clade 3: EV-A76, EV-A89, EV-A90, EV-A91, EV-A119, EV-A121 and EV-A125). Finally, clade 4 contains only NHP EV strains (Clade 4: EV-A122, EV-A123 and EV-A124). Similarly, within EV-D, some types are human-specific (EV-D70, EV-D68 and EV-D94), or NHP-specific (EV-D120), while EV-D111 has been detected in both hosts (Sadeuh-Mba et al., 2019). No unidentified or misidentified NHP derived EV-A -B or -D were found in public databases.

**Figure 4.**
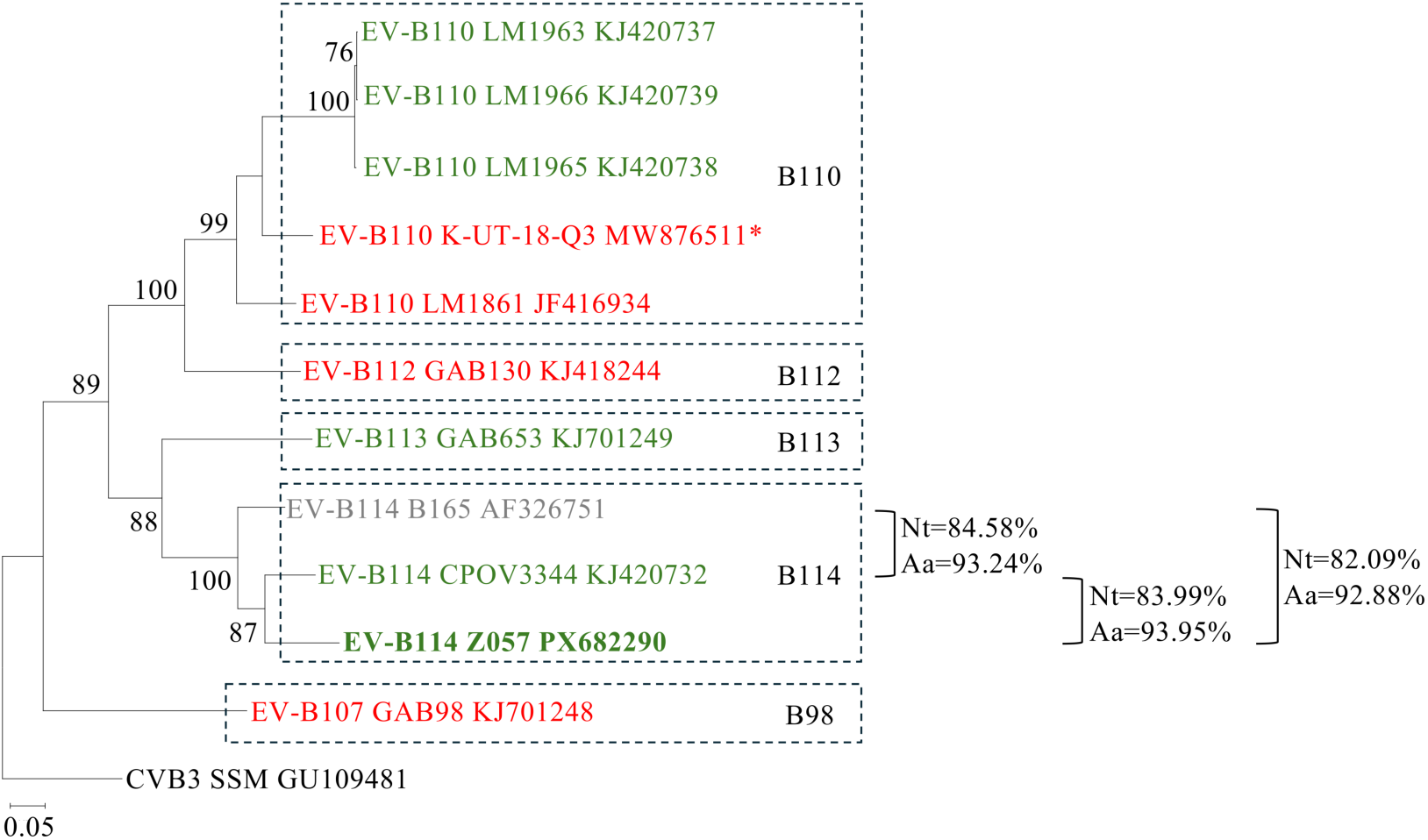
VP1-based phylogenetic tree of enterovirus B species. The maximum-likelihood tree was constructed with all available full NHP EV-B VP1 sequences. The tree is rooted with the EV-B3 SSM strain. The colours of the strain names indicate their host origin: *Mandrillus* (green), *Homo* (black), *Cercopithecus* (grey), and *Pan* (red). Bootstrap values greater than 70% (based on 500 pseudoreplicates) are shown at the corresponding nodes. Percentages of nucleotide (Nt) and amino acid (Aa) identities between EV-B114 strains are indicated. The sequence alignment was performed using the CLC Main Workbench v24, and the phylogenetic tree was constructed with MEGA X. The tree is drawn to scale, with branch lengths measured in the number of substitutions per site.

Notably, 7 sequences spanning the entire main ORF (RCMH3, RCMH6 -both sequenced in this study-, CPML3961, CPML8109, GR2815 and NGR_NHP_1) clustered together in the VP1-based phylogenetic tree and did not align with any known species (Supplementary Figure 1). To determine whether these sequences belong to a single new species, we calculated their pairwise nt and aa identities (Supplementary Figure 6). Our analysis showed that the corresponding viruses met the ICTV criteria for classification as the same species: they share over 70% aa identity in the ORF (85.69%-95.02%) and in the 2C+3CD sequences (97.03%-99.69%), as well as over 60% aa identity in the capsid sequences (69.27%-89.82%). Therefore, we propose the creation of a new species, tentatively named *Enterovirus mbel* (EV-M), after the name of *Pterocarpus soyauxii*, a tree native to the tropical forest in Central African (Albreht et al., 2025), known as *mbel* in various Bantu languages spoken in Central and Southern Cameroon, where the first representatives of this species were sampled (Galley, 1964; Tsala, 1956). Phylogenetic analyses of the VP1 gene revealed several clusters within this new EV-M species (Figure 5). Based on the established thresholds for discriminating EV types (>75% nt identity and >85% aa identity in the VP1 gene) (Brown et al., 2009), we identified 6 distinct virus types (Supplementary Figure 7): EV-M1 (prototype: RCMH6), EV-M2 (prototype: RCMH3), EV-M3 (prototype: CPML3961), EV-M4 (prototype: CPML8109), EV-M5 (prototype: GR2815) and EV-M6 (prototype: NGR_NHP_1). Identity matrices showed that numerous VP1 sequences deposited in GenBank belong to these newly defined EV-M virus types (Supplementary Figure 8 and Figure 5). EV-M1, EV-M2, EV-M4 and EV-M5 virus types gathered sequences that displayed >78.57% nt and >90.62% aa identities between each other. Within the EV-M3 type, all sequences except two (LM1962 and CPOV3334) displayed a nt identity >77.04% with the prototype strain CPML3961, exceeding the 75% threshold. The nt identities of LM1962 and CPOV3334 compared to CPML396 fell slightly below this threshold (73.22% and 74.14%, respectively), but both shared an aa identity >86.0% with the prototype, supporting their classification as EV-M3. Besides the prototype NGR_NHP_1, no additional members of virus type EV-M6 were identified.

**Figure 5.**
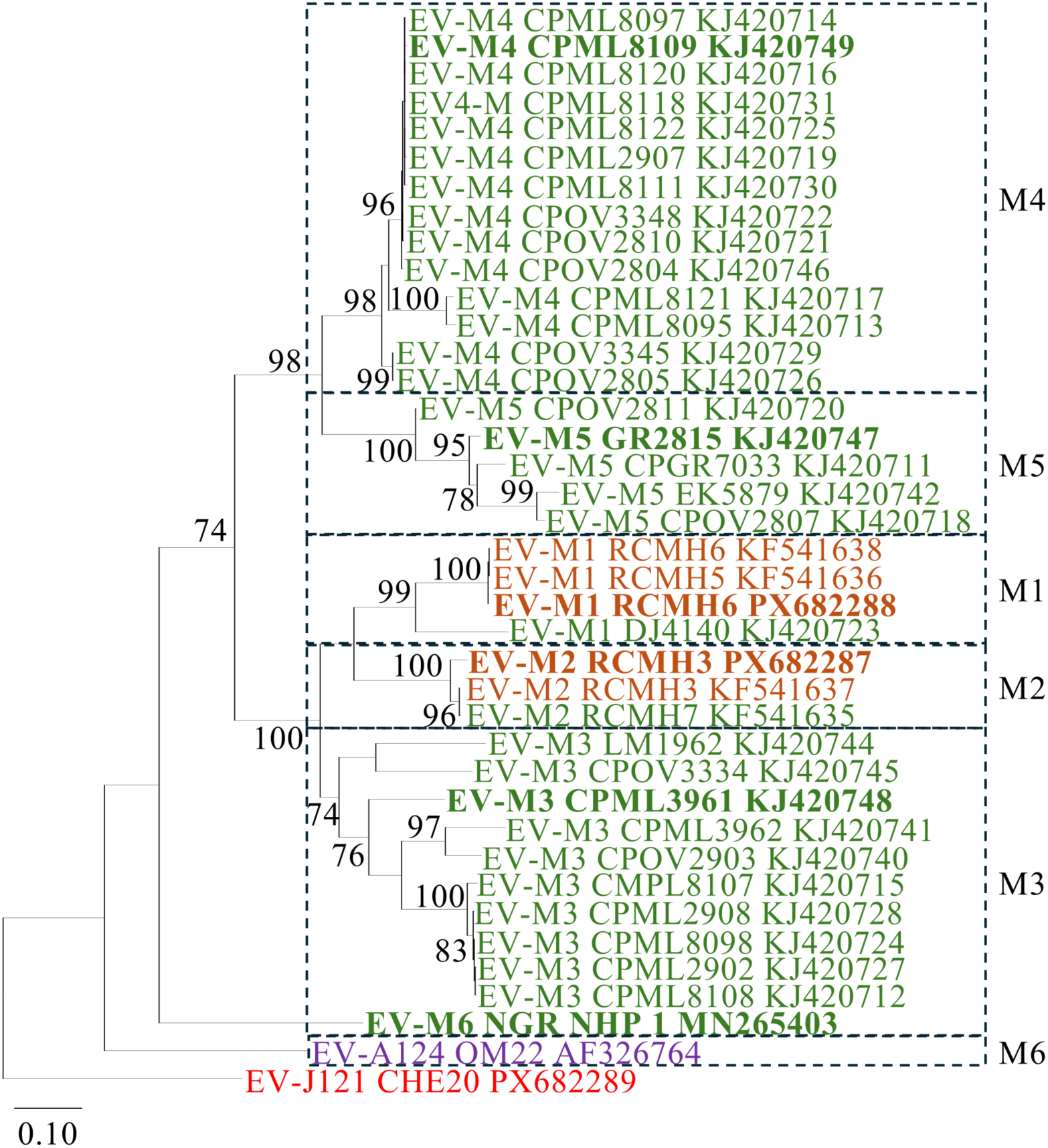
VP1-based phylogenetic tree of the enterovirus M species. The maximum-likelihood tree was constructed with all available full VP1 sequences. VP1 sequences derived from complete genomes are in bold. The colours of the strain names indicate their host origin: *Mandrillus* (green), *Cercocebus* (brown), *Macaca* (purple) and *Pan* (red). The tree is rooted with the EV-J121 CHE20 strain. Bootstrap values greater than 70% (based on 500 pseudoreplicates) are shown at the corresponding nodes. The sequence alignment was performed using the CLC Main Workbench v24, and the phylogenetic tree was constructed with MEGA X. The tree is drawn to scale, with branch lengths measured in the number of substitutions per site.

Shorter VP1 or VP2 sequences, averaging 350 nt in length, have also been published and annotated as belonging to the species EV-J (Mombo et al., 2023, 2021, 2017). However, dendrograms based on these regions revealed that they actually cluster within the EV-M species (Supplementary Figure 9A and 9B). The corresponding sequences are too short to allow accurate typing of the corresponding viruses.

Collectively, these results reveal a complex landscape of NHP EV diversity, including two new species (EV-M and EV-N) and two new EV-J types (EV-J123 and EV-J124). This refined classification will facilitate the proper assignation of NHP EV strains to their respective species and virus type.

### Phylogenetic relationships in the non-structural genomic regions

Recombination events are frequent among co-circulating EVs, generating new viruses with chimeric genomes that may outcompete parental strains if they confer a fitness advantage. Because recombination constantly reshuffles EV genomes, different genomic regions are not subjected to the same extend of selective pressure and do not follow the same evolutionary trajectories. After clarifying the phylogenetic relationships among the NHP derived EVs in the capsid-encoding region, we investigated their evolutionary history by constructing phylogenetic trees considering 2A2B, 2C, 3A3B, 3C, and 3D regions, using all available full-length ORF sequences (Figure 6). As previously reported (Mombo et al., 2017; Oberste et al., 2007; Sadeuh-Mba et al., 2023), striking intra-species phylogenetic incongruences were observed between trees generated from structural (capsid) and non-structural (2A2B, 2C, 3A3B and 3D) regions. Within the EV-B species, the NHP and human derived EVs formed two distinct clades, consistent with the VP1 phylogeny (Figure 6). Of note, all known EV-Ms clustered with the NHP EV-B clade across all non-structural regions. Consequently, EV-Ms and NHP EV-Bs are phylogenetically indistinguishable from each other across the non-structural part of the genome. Similarly, as previously reported (Oberste et al., 2007; Sadeuh-Mba et al., 2023), EV-A125 strain BA13 did not cluster with other EV-As in the non-structural regions downstream the capsid. Our analyses further revealed that BA13 strain clustered with EV-J108 strains in the 2A2B, 2C and 3A3B regions, with the EV-J103 strains in the 3C gene and with the members of the new species EV-N in the 3D gene (Figure 6). Finally, the species EV-D, EV-H, and EV-L exhibited complex phylogenetic relationships. Indeed, they form a single group in the 2A2B region, but segregated into three distinct branches in the 3A3B and 3C based trees. The other regions showed specific pairwise associations: EV-D grouped with EV-H and EV-L in the 2C region and the 3D region, respectively.

**Figure 6.**
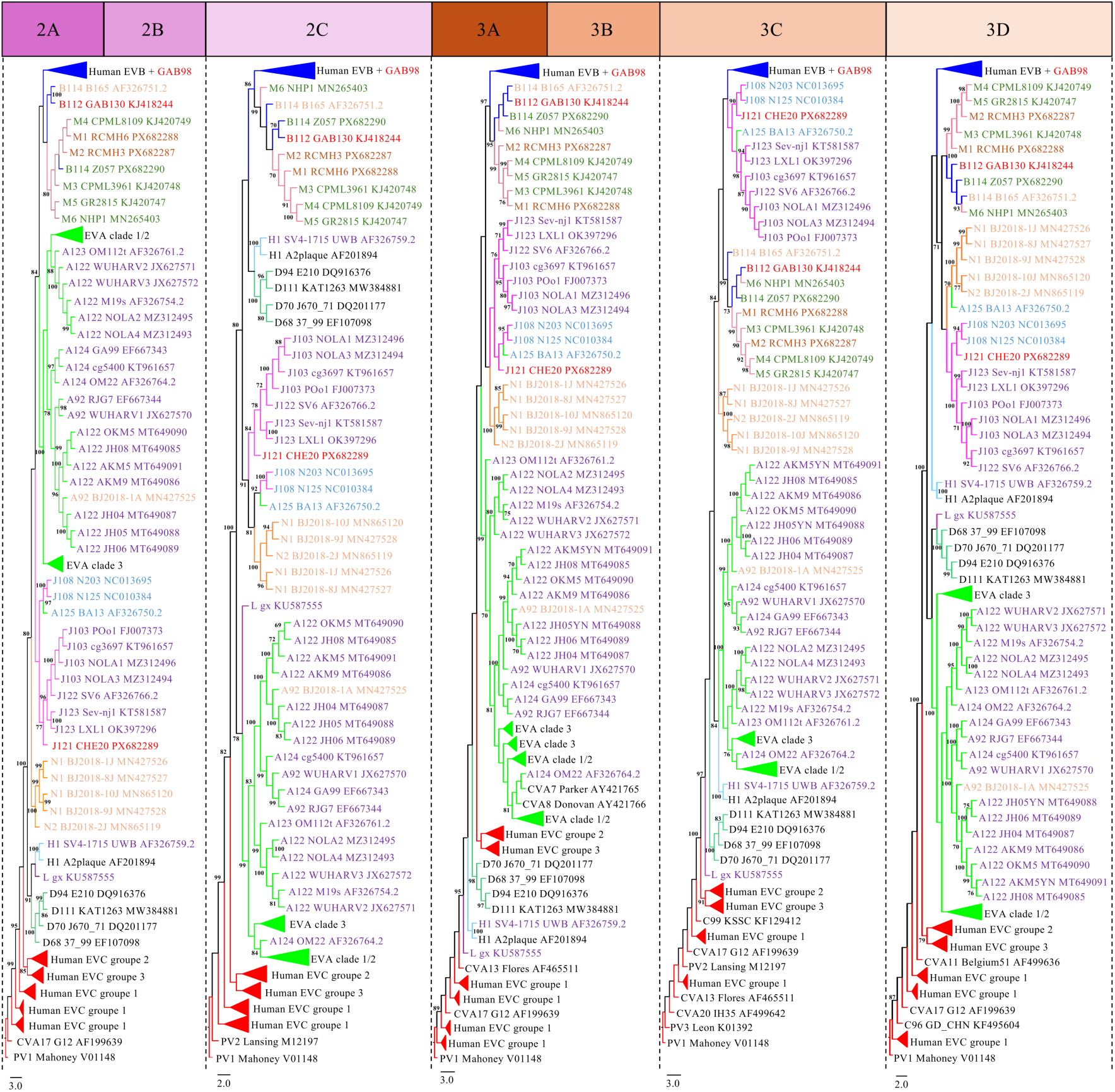
Phylogenetic analysis of non-structural genes derived from complete genomes of NHP and representative human EVs. The maximum-likelihood trees were constructed from the 2A2B, 2C, 3A3B, 3C and 3D regions of all complete NHP and representative human EV genomes. Branches are coloured according to EV species: EV-A (light green), EV-B (dark blue), EV-C (red), EV-D (dark green), EV-H (light blue), EV-J (pink), EV-L (purple), EV-M (brown) and EV-N (orange). The colours of the strain names indicate their host origin: *Macaca* (purple), *Pan* (red), *Papio* (light blue), *Chlorocebus* (orange), *Mandrillus* (green), *Cercocebus* (brown) and *Homo* (black). The trees are rooted with the PV1 Mahoney strain (V01148). Bootstrap values greater than 70% (based on 500 pseudoreplicates) are shown at the corresponding nodes. The sequence alignments were performed using the CLC Main Workbench v24, the phylogenetic trees were constructed with MEGA X, and the visualizations were generated with FigTree. The trees are drawn to scale, with branch lengths measured in the number of substitutions per site.

### Comparison of genomic signatures of human and NHP derived EVs

To our knowledge, no comparative genomic analysis of human and NHP derived EV species has been published so far. A previous study, limited to EV-A species, revealed that clade 3 viruses displayed distinct relative dinucleotide ratios and codon usage biases compared to human EV-As of clades 1 and 2 (Zeng et al., 2022), suggesting that specific genomic signatures could be associated with the virus’ natural history and, particularly its host range. To determine whether NHP derived EVs from different host species shared unique genomic traits, we compared GC content (GC%), dinucleotide ratio, and codon usage (ENC and RSCU analysis) between human and NHP derived EVs. To minimize potential bias in these analyses, we included, where possible, at least the prototype strain and two additional genomes sampled from different countries for each virus type.

The GC% ranged from 41% to 49% across all species (Figure 7A). Interestingly, the GC% strongly correlated with the host origin rather than EV species classification, with all NHP virus types featuring a GC% <47%. In particular, NHP-specific EV species (EV-J, EV-H, EV-L, EV-M and EV-N) display low GC%, ranging from 43% to 46.4%, as did the few known EV-D virus types (41.0%-43.5%). Strikingly, virus types segregated with respect to their host of origin within the EV-A and EV-B species: all EV-A virus types of clades 1 and 2, originating from humans, displayed GC%>47%, while NHP-associated clades 3 and 4 had GC%<47% (Figure 7B). A similar clustering was found within EV-B species, with all human derived EV types featuring a GC% >47%, while all NHP derived EV types had a GC% <45% (Figure 7C). Notably, the GC % of EV-C virus types, which are restricted to human hosts, mirrored their phylogenetic relationships: respiratory virus types (EV-C104, EV-C105, EV-C109, EV-C117 and EV-C118) exhibited a high GC % (48–48.5%), cultivable enteric virus types (PV1, PV2, PV3, CVA11, CVA13, CVA17, CVA20, CVA21, CVA24, EV-C95, EV-C96, EV-C99 and EV-C102) displayed an intermediate GC % (45–46.5 %), while non-cultivable viruses (CVA1, CVA19, CVA22, EV-C113, EV-C116 and EV-C119) showed a low GC % (43-44 %) (Figure 7D).

**Figure 7.**
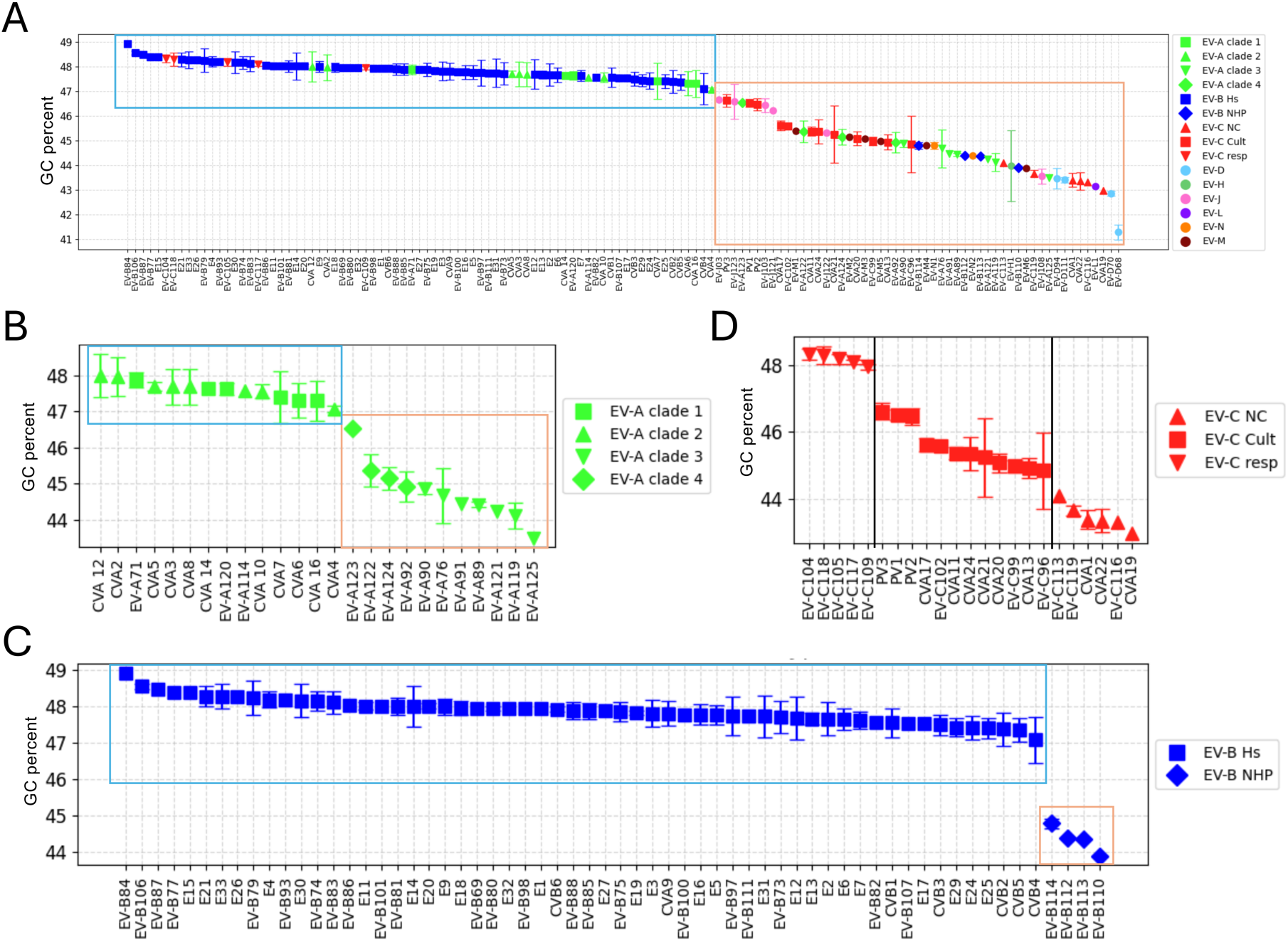
Distribution of GC content across enterovirus species. GC content distribution for all EV species **(A)**, EV-A species **(B)**, EV-B species **(C)**, and EV-C species **(D)** are plotted. Blue boxes indicate clusters of human derived EV while brown boxes indicate clusters of mainly NHP derived EVs. When possible, three sequences per type were included in the analysis and standard deviations are plotted. Abbreviations: Cult, cultivable; NC, non-cultivable; Resp, respiratory; Hs, *Homo sapiens*; NHP, non-human primate.

A previous study revealed that EV-As clade 3 viruses display lower CpG and UpA dinucleotide frequencies compared to clades 1 and 2 (Zeng et al., 2022). To determine whether NHP derived EV-As of clade 4 share this trait, we analyzed dinucleotide ratios of EV-A strains representative of the different clades (Figure 8). Our analysis revealed that EV-A viruses belonging to clade 4 had the same low CpG ratio as clade 3 EV-As (Figure 8B). The same observation was made within EV-B species, with NHP derived EV types displaying a CpG ratio substantially lower than that of human derived EV types (NHP EVs<0.42 vs Human EVs>0.56) (Figure 8C). With the exception of EV-H, a low CpG ratio seems to be a common characteristic of NHP derived EVs. Indeed, members of the EV-J, EV-L, EV-M and EV-N species featured CpG ratios<0.5 and comparable to those of NHP derived EV-A and EV-B species (Figure 8A). It is noteworthy that all EV-D virus types had CpG ratio<0.35, irrespective of their host of origin.

**Figure 8.**
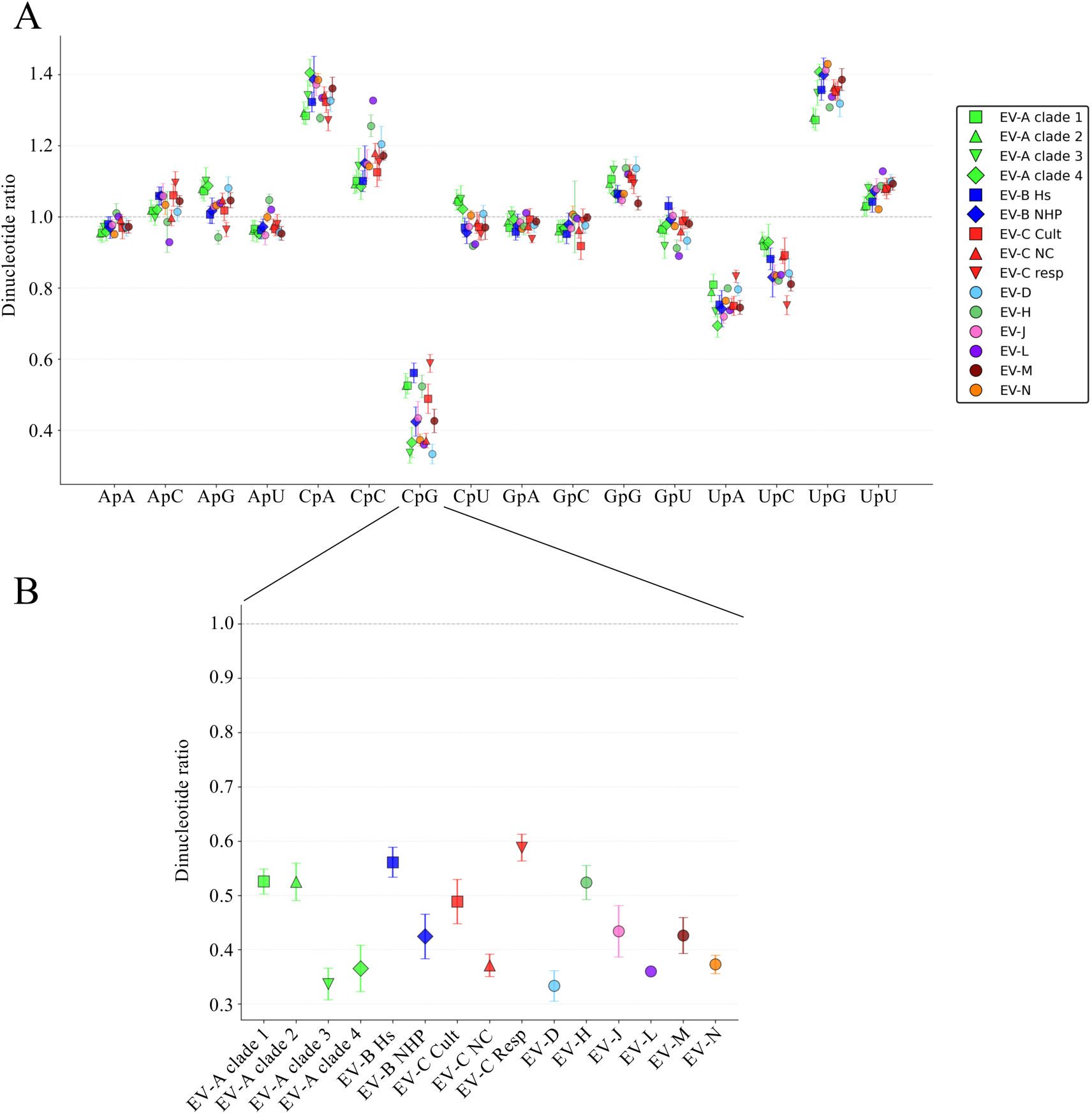
Dinucleotide ratio analysis of human and non-human primate enterovirus genomes. **(A)** All dinucleotide ratios are shown. **(B)** The CpG dinucleotide ratio is specifically highlighted for each EV species. Abbreviations: Cult, cultivable; NC, non-cultivable; Resp, respiratory; Hs, *Homo sapiens*; NHP, non-human primate.

We also investigated codon usage bias in human and NHP derived EVs. The effective number of codons (ENC), which ranges from 20 (strong bias, with one codon used for each amino-acid) to 61 (no bias, even use of each codon) (Wright, 1990), revealed distinct patterns. Human EV-As (clades 1 and 2), EV-Bs, respiratory EV-Cs, and NHP derived EV-Hs had ENC greater than 54 (Figure 9A), while clade 3, NHP derived EV-As, NHP derived EV-Bs, cultivable and non-cultivable EV-Cs, EV-Ds, EV-Js and EV-Ms featured ENC values between 50 and 54. The sole known EV-L had an ENC value of 48. Since the GC% at the third codon position (GC3) could introduce a bias in the ENC analysis, we compared the ENC values for each EV species with the expected ENC based on the respective GC3 percentage (Butt et al., 2016) (Supplementary Figure 10). All calculated ENC values fell below the expected curve, confirming that GC3 content did not account for the variations across virus group.

**Figure 9.**
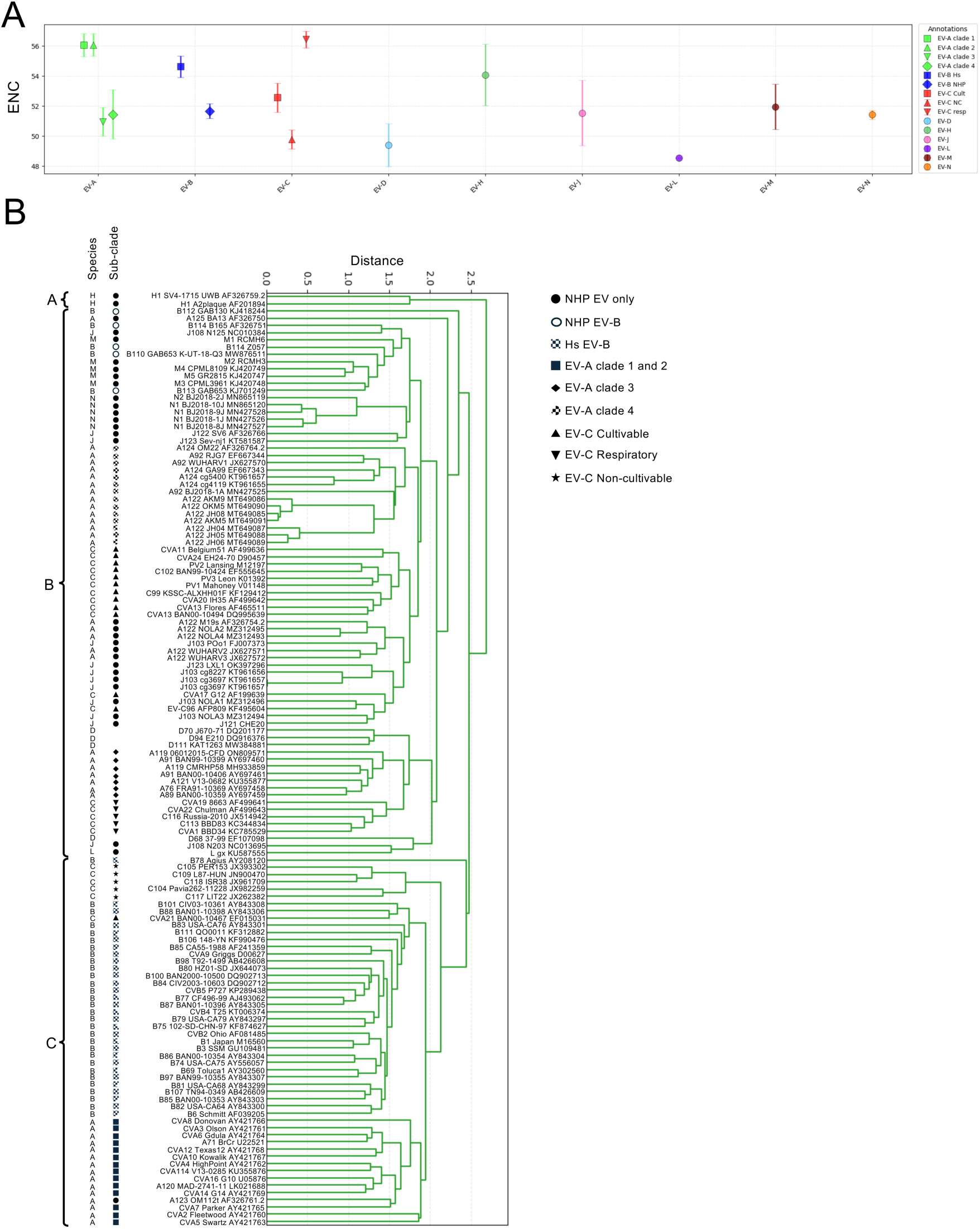
Codon usage bias analysis of enterovirus species. **(A)** Effective number of codons (ENC) values for all NHP EVs and representative human EVs are plotted. When possible, three sequences per type were included in the analysis. **(B)** Relative synonymous codon usage (RSCU) values for all NHP EVs and representative human EVs were summarized in a cladogram. The three main groups separated by the cladogram are labeled A, B and C. Abbreviations: Hs, *Homo sapiens*; NHP, non-human primate.

We also analyzed the Relative Synonymous Codon Usage (RSCU), which quantifies how frequently each synonymous codon is used in a gene or genome relative to the expected usage. The results, summarized in a cladogram, grouped EV types into three main clusters (Figure 9B). Group A contains only the EV-H1 type. Group B includes all other NHP-specific EV types (except EV-A123), whichever their species, all human cultivable EV-Cs, all EV-Ds and EV-A clades 3 and 4. Group C contains all other human EV-Cs and EV-A123. These results suggest that codon-usage bias varies markedly among EV groups, yet it only partially mirrors the underlying phylogenetic relationships among them.

Collectively, these results reveal that dinucleotide ratios, ENC and RSCU values vary among NHP EV species. Remarkably, GC% mirrors phylogenetic relationships between EVs and could serve as a predictive marker of human versus NHP origin of EVs.

## DISCUSSION

For decades, no standardized nomenclature existed for NHP derived EVs. In the 1980s, it was proposed to designate these viruses under the common name, *Simian enterovirus*, followed by a number corresponding to a given serotype (Kalter, 1982). For a long time, NHP derived EVs were not assigned to specific species. In the 2000s and 2010s, the generalization of genetic sequencing brought about major changes in EV classification, as phylogenetic relationships replaced phenotypic criteria to delineate EV types and species. Based on their genetic relationships, NHP derived EVs were classified into different species, some of which also included human derived EVs. By considering all NHP EV genome sequences available in public databases and integrating our two new full-genome sequences, our analyses confirmed the existence of a new species (Figure 5 and Supplementary Figure 6) that had been previously suggested by two independent studies (Nguyen et al., 2014; Sadeuh-Mba et al., 2014). We found that this new EV species gathered viruses sampled in three countries, Gabon, Cameroon, and Nigeria (Mombo et al., 2023, 2021, 2017; Nguyen et al., 2014; Sadeuh-Mba et al., 2023). This geographic distribution justifies the name proposed for this species, *Enterovirus mbel*, which refers to an endemic tree of this region, called African padauk in English (Albreht et al., 2025). We also identified another new EV species, comprising a few viruses sampled in China previously misidentified as EV-Js (Li et al., 2020) (Figure 3 and Supplementary Figure 5). Since this species has only been documented in China, we propose naming it *Enterovirus nao,* after the Chinese word referring to a monkey in a poem written 2,500-3,000 years ago (Couvreur, 1896; Loewe, 1993). However, it is important to note that the genuine geographic area of circulation of these two new EV species remain to be documented, especially for *E. nao* which was detected in captive animals.

The discovery of the uORF in the genome of some EVs a few years ago constituted a drastic change in our understanding of their biology. This uORF was experimentally shown to encode a non-essential protein that enhances virus replication in gut cells (Lulla et al., 2019). Interestingly, the putative uORFs identified in the genomes of EV-B114 Z057, EV-M1 RCMH6, and EV-M3 RCMH3 are respectively located 134 nt, 133 nt, and 136 nt upstream of the AUG start codon of the main ORF. These distances fall below the cutoff set by Lulla et al. for screening for uORF of EV genomic sequences (Lulla et al., 2019). Although functional validation of these candidate uORFs is still lacking, our findings suggest that the prevalence of uORFs across EVs may have been underestimated and warrants re-evaluation using a more permissive distance threshold.

The phylogenetic analyses performed on the non-structural part of the genome further confirmed the close genetic relationships between NHP EVs recovered from different species, as previously documented (Figure 6) (Mombo et al., 2017; Nguyen et al., 2014; Oberste et al., 2007; Sadeuh-Mba et al., 2014). This close genetic relatedness could be explained by convergent evolution, which may have driven EV genomes from different species to become phylogenetically more similar as a consequence of their replication within NHP hosts. However, it is unlikely that convergent evolution could uniformly affect the entire non-structural region of the genome, as it would be the case for NHP derived EV-Bs and EV-Ms that cluster together across this region. Furthermore, convergent evolution among NHP derived EVs is not supported by the fact that most known NHP derived EV-As do not cluster with other NHP derived EVs, except for EV-A125 strain BA13, which is phylogenetically close to EV-Js downstream the capsid. If convergent evolution was shaping the evolution of EV-A125, it would be expected to similarly affect other NHP derived EV-As, which seems unlikely. Therefore, these phylogenetic incongruences found in this study are probably due to recombinations, a well-established driver of EV evolution (Muslin et al., 2019).

While interspecies recombination has been observed in the 5’UTR (Boros et al., 2012; Muslin et al., 2015; Santti et al., 1999), it has never been documented downstream of the capsid-encoding region among human derived EVs. Even intraspecies recombination is limited to specific subsets of EV types that can recombine with one another but not with members of other subsets. For instance, EV-As of clade 3 do not exchange genetic sequences with EV-As of clades 1 and 2 (Oberste et al., 2005; Sadeuh-Mba et al., 2014). Similarly, respiratory EV-Cs and non-cultivable EV-Cs do not recombine with each other or with cultivable EV-Cs (Brouwer et al., 2020). Several factors could explain the apparent inability of some EVs to generate recombinant viruses, such as functional incompatibilities between genomic regions from distant phylogenetic groups, lack of fitness advantages conferred to the recombinant viruses, or low frequency of co-infection due to different host and tissue tropisms. Our analysis provided substantial evidence to the existence of interspecies recombination between some NHP derived EV species, particularly between EV-Ms and EV-Bs. These findings change our understanding of recombination between EVs, which appear to be capable of producing larger leaps than previously believed. This raise fundamental questions about the past and future evolutionary path of these viruses. For instance, it is unknown whether NHP derived EV-Bs acqiured non-structural genes from EV-Ms, or vice-versa, unless both inherited these genes from a yet-unidentified common source. The ability of EV-Ms to recombine with human derived EV-Bs and the potential for such inter-species macro-evolutionary events to produce new human-adapted EV-Ms are also difficult to predict.

Previous studies have shown that genomes of clade 3 EV-As, which were detected in both NHPs and humans, had a lower CpG frequency than human specific EV-As and featured a specific codon usage bias that impairs their expression of these EV-A genomes in human cells (Zeng et al., 2022). These observations led us to hypothesize that some genetic characteristics shared by NHP derived EVs may play a role in host adaptation. Nucleotide composition has been previously used to predict the host of novel picorna-like viruses (Kapoor et al., 2010). While our analysis of dinucleotide frequency and codon usage bias across the various EV species circulating in humans and in NHPs did not reveal a clear segregation between human and NHP derived viruses (Figure 9), our results identified a relatively low GC% appears as a signature shared by virus types circulating among NHPs, regardless of their virus species (Figure 7). Unfortunately, the lack of data precluded robust statistical analysis, as some EV types are represented by hundreds of sequences while others, such as EV-L species, are represented by only one or two sequences. Based on genetic and epidemiological data, we and others have previously postulated that EV-Ds known in humans may have recently emerged from zoonotic sources, presumably NHPs (Kono et al., 1981; Miyamura et al., 1986; Sadeuh-Mba et al., 2019). The low GC% values displayed by all EV-D virus types, irrespective of their tissue tropism or host of origin, are consistent with that hypothesis. The relative relatedness of EV-Ds and NHP-specific EV species in the non-structural part of the genome further supporting the hypothesis of a zoonotic origin of EV-Ds. However, host adaptation is not the only driver of GC%, as illustrated by EV-C virus types, which display a broad range of GC% values despite all circulating in humans (Figure 7). In addition, the GC% of human and NHP EV strains falls within the broad range observed in the human transcriptome (average 48%, varying from <40% to >60% among mRNAs) (Courel et al., 2019; Piovesan et al., 2019). Unfortunately, the GC% of transcriptomes of NHP infected with EVs remains uncharacterized to our knowledge. The GC% overlap between EVs and human transcriptome is insufficient to conclude host-species adaptation, and the lack of data prevents comparative analysis that could elucidate potential host-driven GC% adaptation in NHP EV strains. The genomic signature result from a dynamic equilibrium influenced by many factors, of which polymerase fidelity, RNA secondary structure requirements, host translational preferences, immune pressure, and genetic recombination. Furthermore, identifying genomic signatures that reliably distinguish NHP derived EVs from human derived EVs is currently limited by the paucity of fully sequenced strains, our limited knowledge of the natural history of these viruses, and particularly the uncertainty surrounding their host tropisms.

## ACKNOWLEDGEMENT

The authors are indebted to Dr. WEI Yu, who shared her knowledge of ancient Chinese poetry, and Dr. Nick J. Knowles, for his feedback regarding enterovirus phylogeny. This work has used the computational and storage services (Maestro cluster) provided by the IT department at Institut Pasteur, Paris.

## AUTHORS CONTRIBUTIONS

MB and SASM conceived the study. SASM, PCdC, MP, MLJ, MCEZ, AB and ESL generated the sequencing data. CA, ESL, and MB performed the analyses. CA, NJ and MB wrote the article, which was approved by all co-authors.

---

**Supplementary Table 1.** Predicted polyprotein cleavage sites of enterovirus Z057, CHE20, RCMH3, and RCMH6. Cleavage site predictions are based on multiple sequence alignment with the annotated poliovirus type 1 Mahoney reference strain (V01148.1). The analysis was performed using CLC Main Workbench v24 software.

| Strains | VP4/VP2 | VP2/VP3 | VP3/VP1 | VP1/2A | 2A/2B | 2B/2C | 2C/3A | 3A/3B | 3B/3C | 3C/3D |
| --- | --- | --- | --- | --- | --- | --- | --- | --- | --- | --- |
| **CHE20** | LK/SP | TQ/GL | LQ/SP | TA/GA | EQ/GI | KQ/AD | FQ/GP | FQ/GA | VQ/GP | EQ/GE |
| **Z057** | LN/SP | PQ/GL | LQ/SP | TY/GA | EQ/GV | RQ/AD | FQ/GP | FQ/GA | VQ/GP | EQ/GE |
| **RCMH3** | LK/SP | TQ/GL | LQ/GD | NT/GA | EQ/GV | RQ/AD | FQ/GP | FQ/GA | VQ/GP | EQ/GE |
| **RCMH6** | LK/SP | AQ/GL | LQ/GD | TT/GA | EQ/GV | RQ/AD | FQ/GP | FQ/GA | VQ/GP | EQ/GE |

**Supplementary Figure 1.**
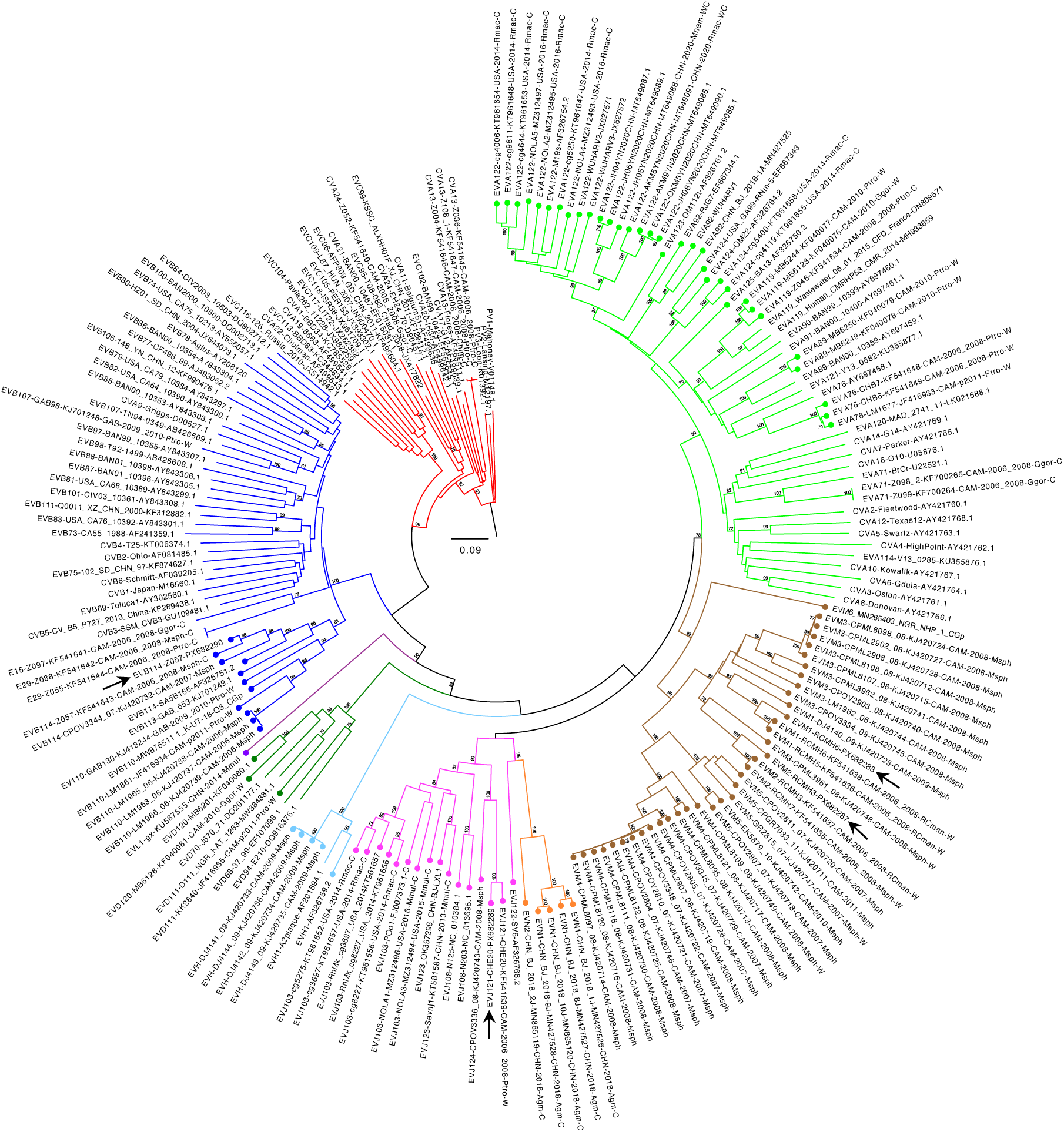
Phylogenetic analysis of complete VP1 sequences from non-human primate and representative human enteroviruses. This phylogenetic tree is the same as in the Figure 2, with the name of each EV type indicated. For NHP EV, the following nomenclature is used: Species-Type-Strain name-Accession number-Country-Year-Host species-Condition of life of host. For prototypic EVs, the following nomenclature is used: Species-Type-Strain name-Accession Number. Abbreviations: CAM, Cameroon; CHN, China; USA, United States of America; GAB, Gabon; NGR, Nigeria; Msph, *Mandrillus sphinx*; Mleu, *Mandrillus leucophaeus*; Ctor, *Cercocebus torquatus*; Caet, *Chlorocebus aethiops*; Ptro, *Pan troglodytes*; Ggor, *Gorilla gorilla*; Mmul, *Macaca mulatta*; W, wild; C, captive. The arrows indicate new genomes sequenced in our study (Figure 1). The tree is drawn to scale, with branch lengths measured in the number of substitutions per site.

**Supplementary Figure 2.**
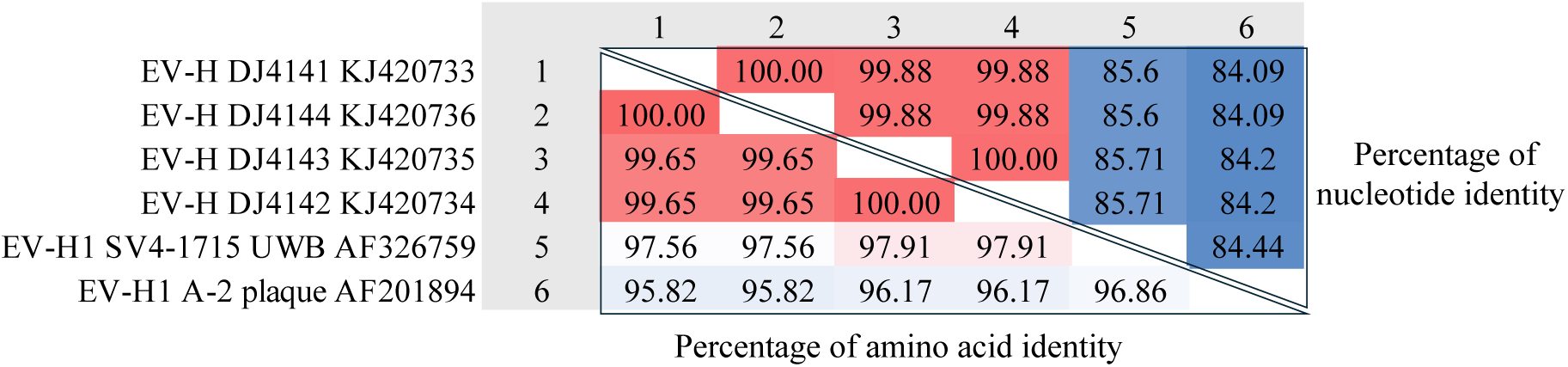
Type characterization of the enterovirus H species. Nucleotide and amino acid identity matrices of EV-H complete VP1 sequences were calculated using CLC Main Workbench v24.

**Supplementary Figure 3.**
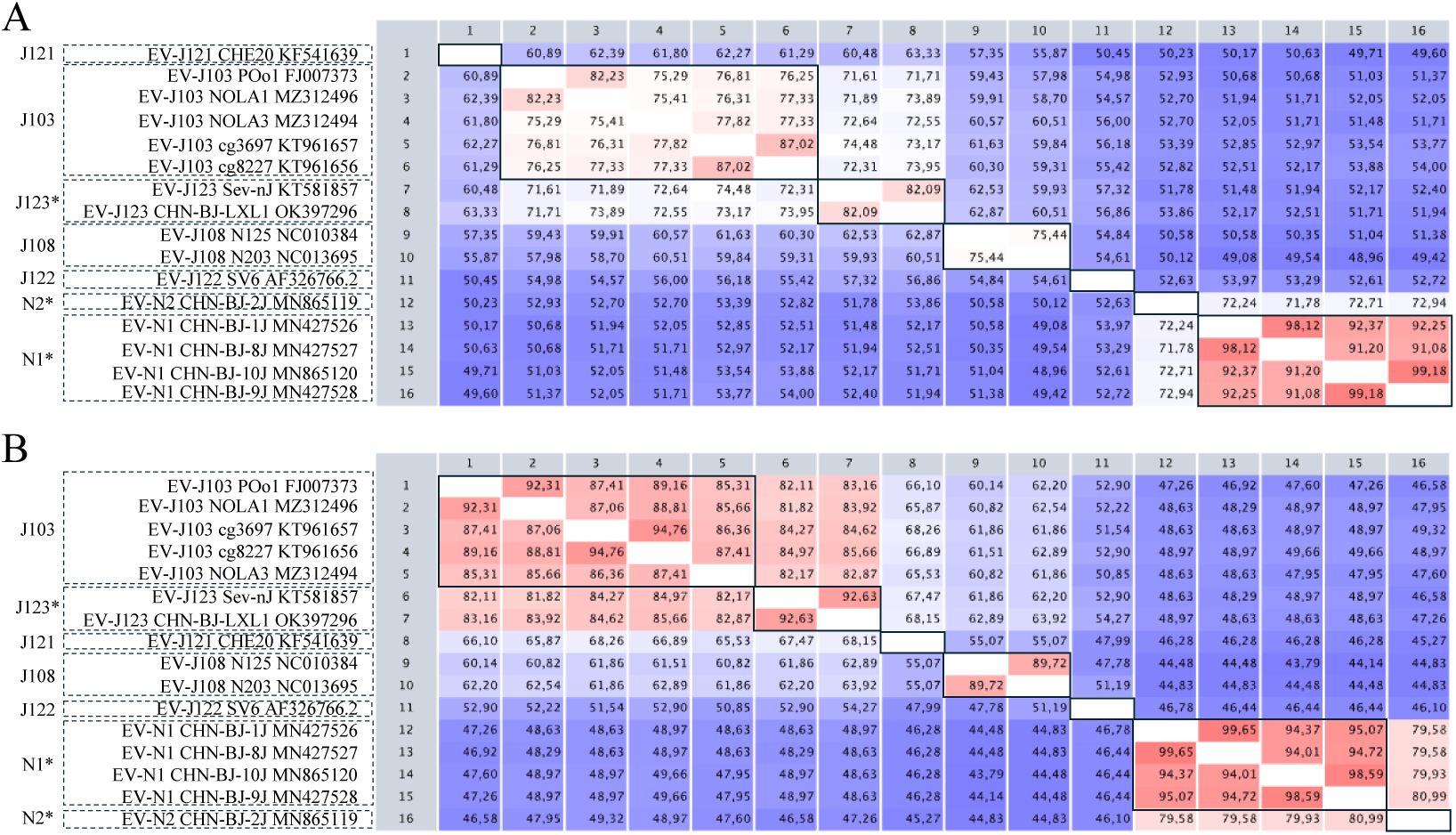
Type characterization of enterovirus J strains based on complete VP1 sequences. Nucleotide **(A)** and amino acid **(B)** identity matrices of VP1 sequences are shown. Translation of nucleotide sequences, multiple sequence alignments, and identity matrices were performed using CLC Main Workbench v24. Putative novel type assignments are indicated by an asterisk (*).

**Supplementary Figure 4.**
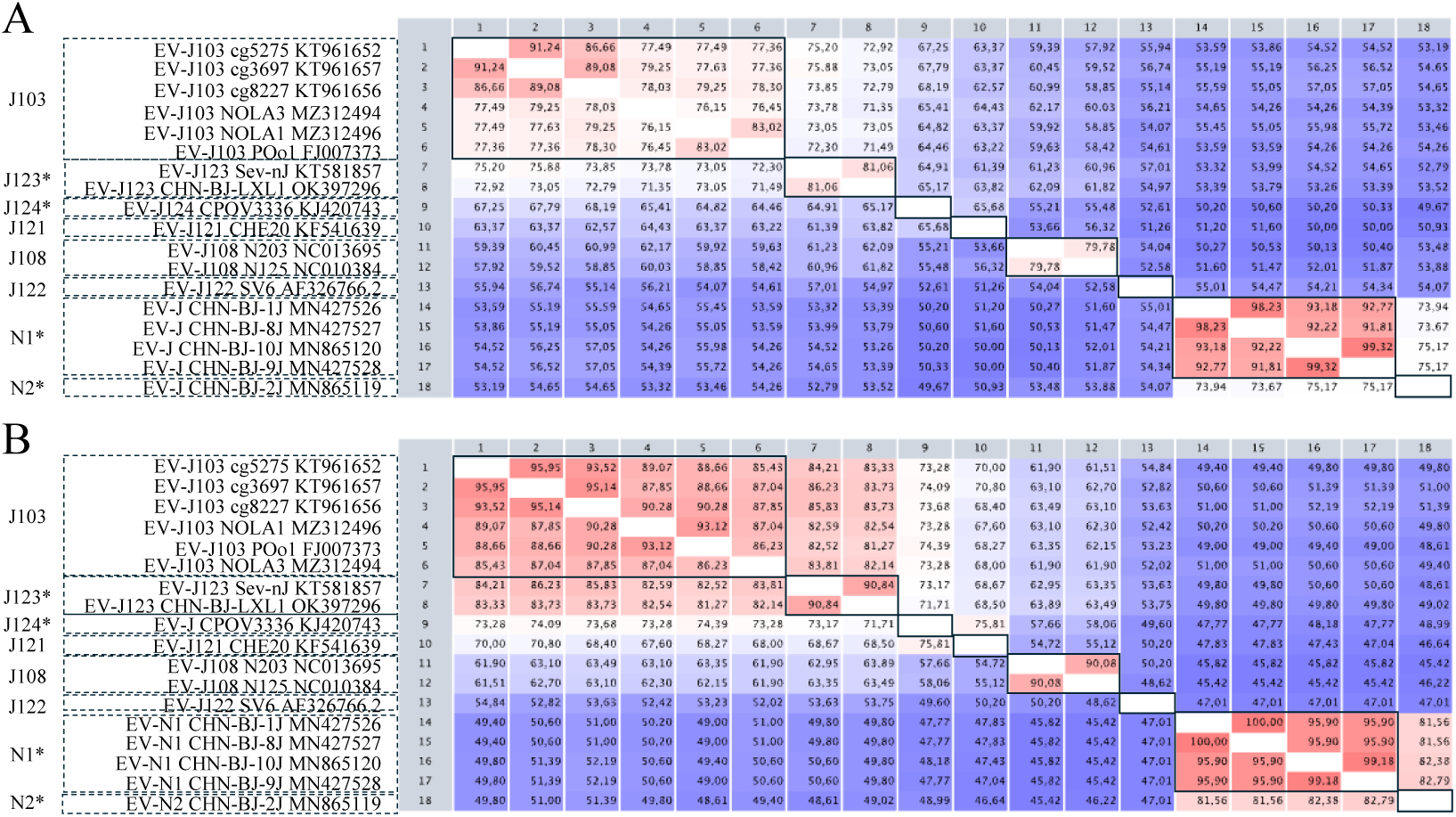
Type characterization of enterovirus J strains based on partial VP1 sequences. Nucleotide **(A)** and amino acid **(B)** identity matrices of partial VP1 sequences are shown. Translation of nucleotide sequences, multiple sequence alignments, and identity matrices were performed using CLC Main Workbench 24. Putative novel type assignments are indicated by an asterisk (*).

**Supplementary Figure 5.**
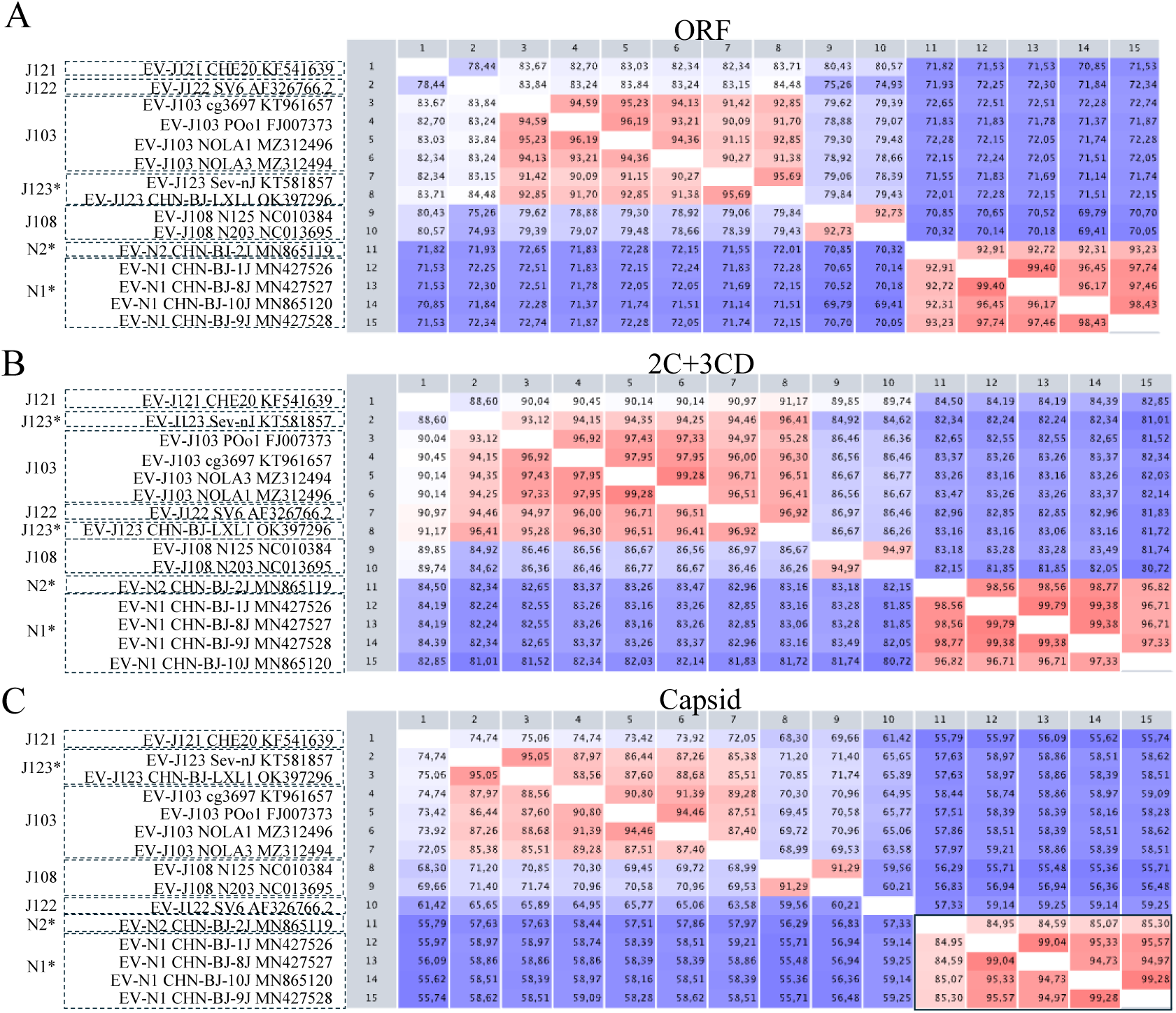
Species characterization of enterovirus J strains. Amino acid identity matrices of **(A)** the complete ORF, **(B)** the 2C+3CD and **(C)** the capsid sequences are shown. Translation of nucleotide sequences, multiple sequence alignments, and identity matrices were performed using CLC Main Workbench v24. Putative novel type assignments are indicated by an asterisk (*).

**Supplementary Figure 6.**
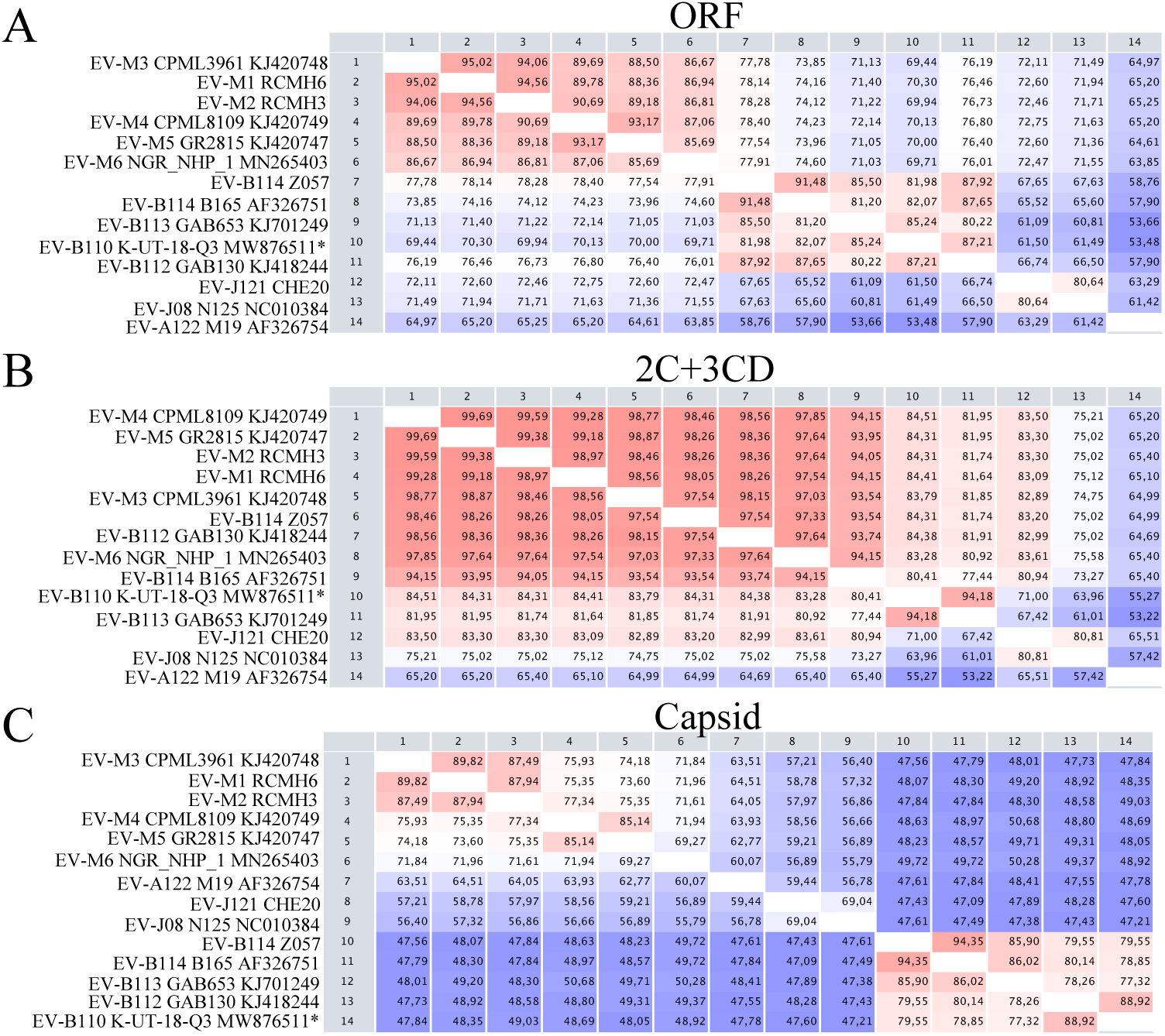
Species characterization of enterovirus M strains. Amino acid identity matrices of **(A)** the complete ORF, **(B)** the 2C+3CD and **(C)** the capsid sequences are shown. Translation of nucleotide sequences, multiple sequence alignments, and identity matrices were performed using CLC Main Workbench v24. The asterisk (*) indicates the sequence where the last 399 nt are missing in the 3D gene.

**Supplementary Figure 7.**
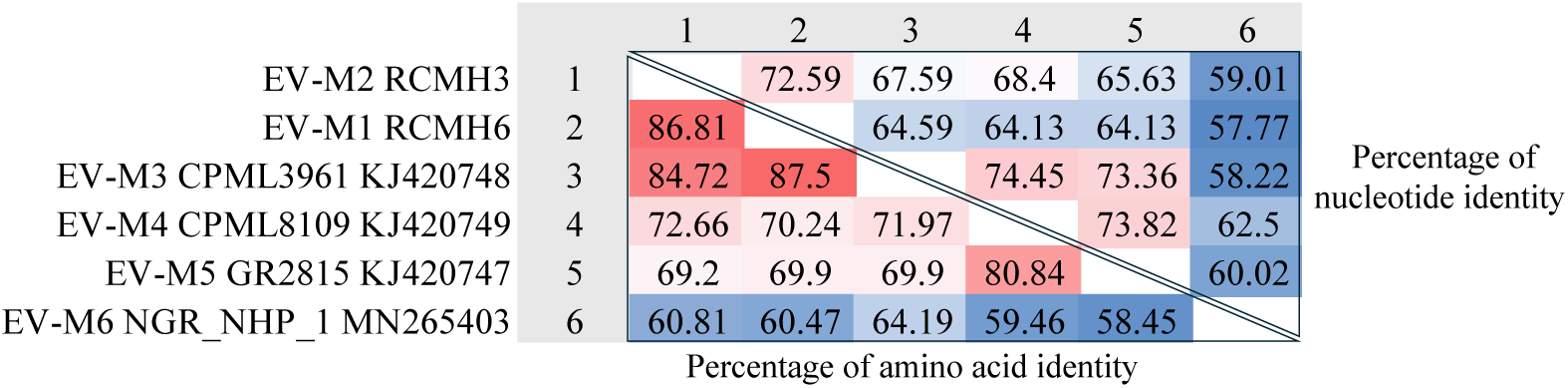
Type characterization of enterovirus M strains based on complete VP1 sequences. Nucleotide and amino acid identity matrices of EV-M VP1 sequences derived from complete genomes were calculated using CLC Main Workbench v24.

**Supplementary Figure 8.**
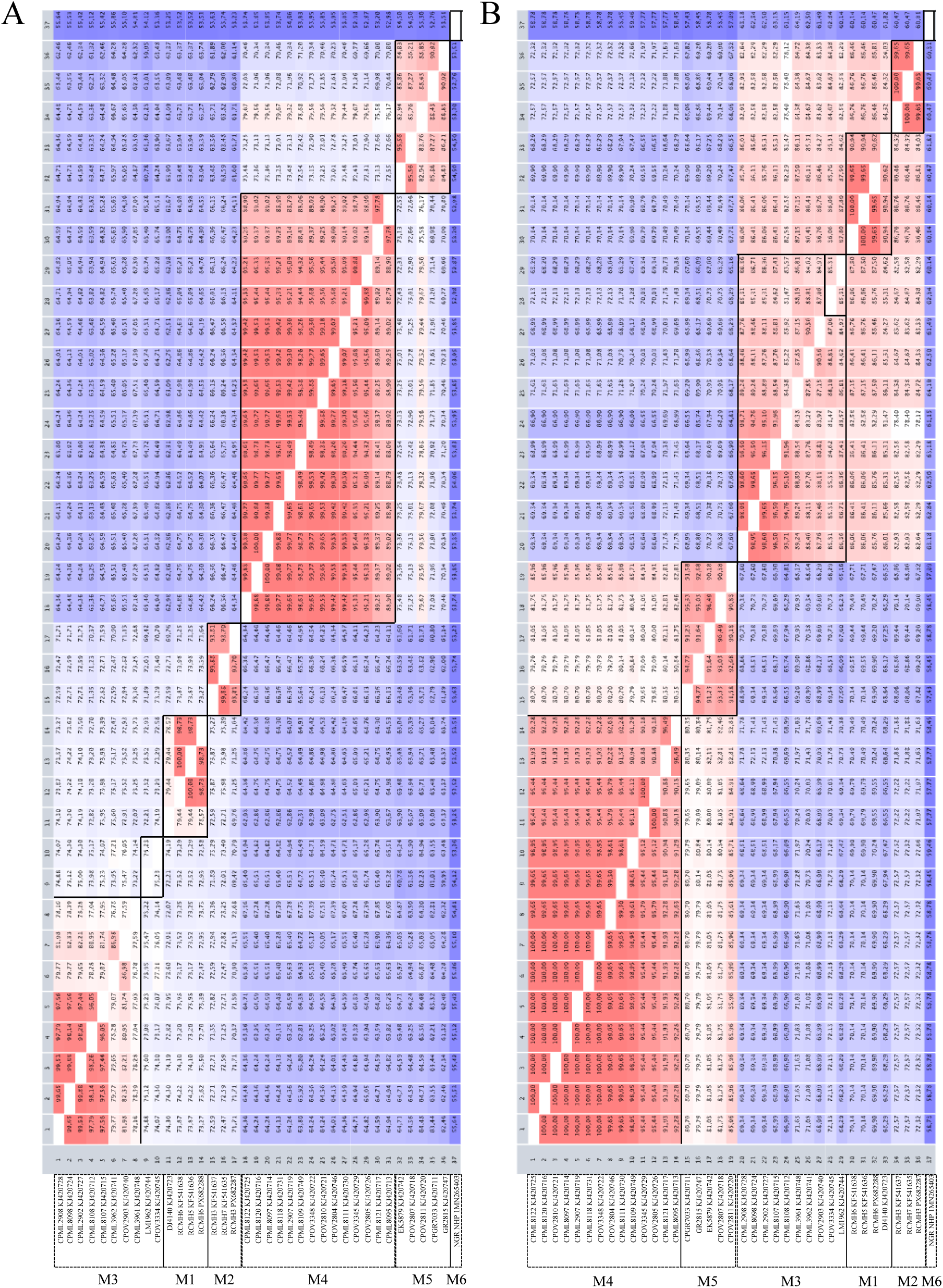
Type characterization of enterovirus M strains based on all available VP1 sequences. Nucleotide **(A)** and amino acid **(B)** identity matrices of VP1 sequences are shown. Translation of nucleotide sequences, multiple sequence alignments, and identity matrices were performed using CLC Main Workbench v24.

**Supplementary Figure 9.**
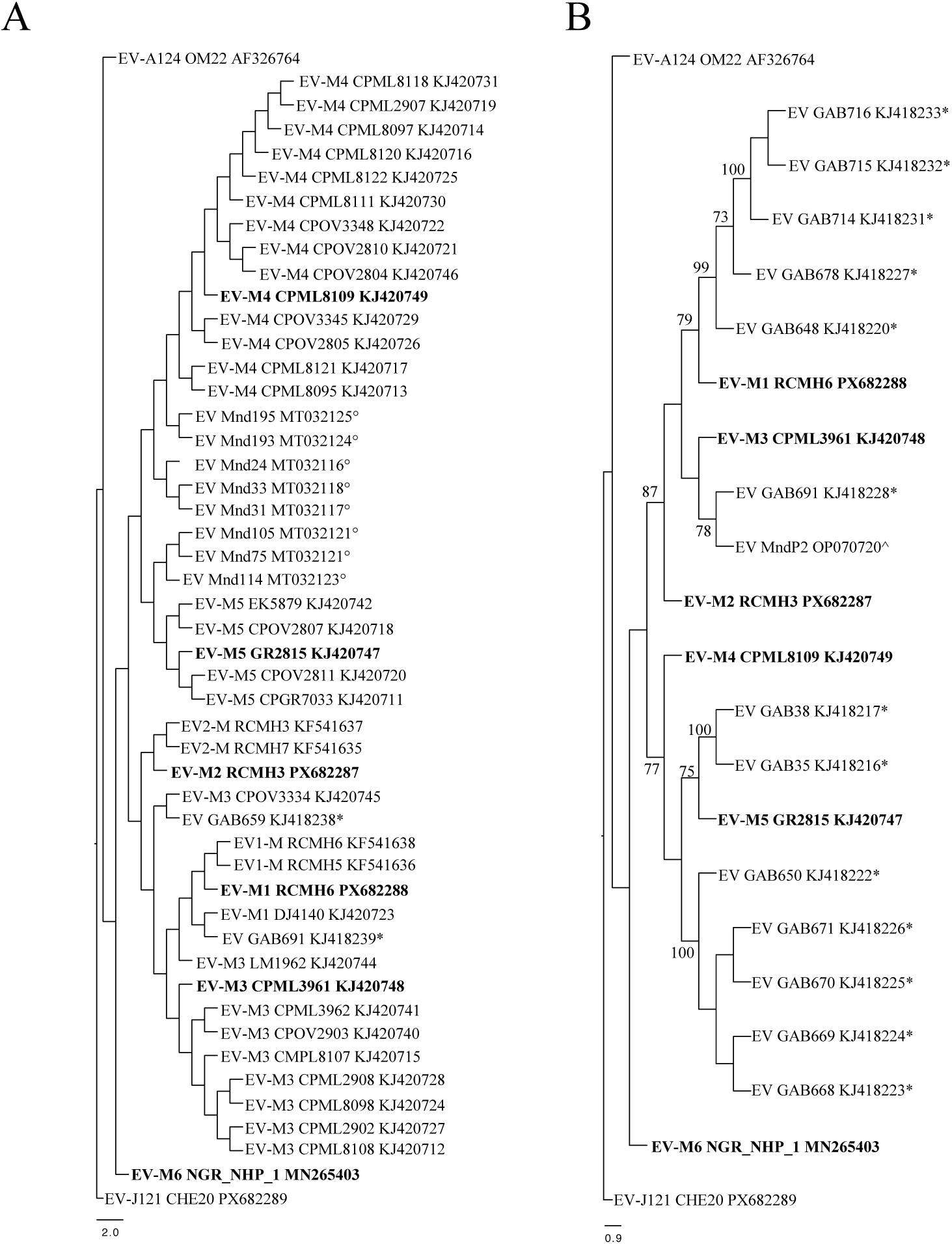
Phylogenetic analysis of enterovirus M strains based on partial VP1 and VP2 sequences. Maximum likelihood phylogenetic trees based on **(A)** VP1 (average length: 354 nucleotides) and **(B)** VP2 (average length: 320 nucleotides) partial sequences are shown. Sequence alignments were performed using CLC Main Workbench v24 and phylogenetic trees were constructed with MEGA X (version 10.2.6). The trees are drawn to scale, with branch lengths measured in the number of substitutions per site. Complete genomes are in bold and symbols indicate references of published sequences: ^, Mombo et al. 2023; *, Mombo et al. 2017; °, Mombo et al. 2021.

**Supplementary Figure 10.**
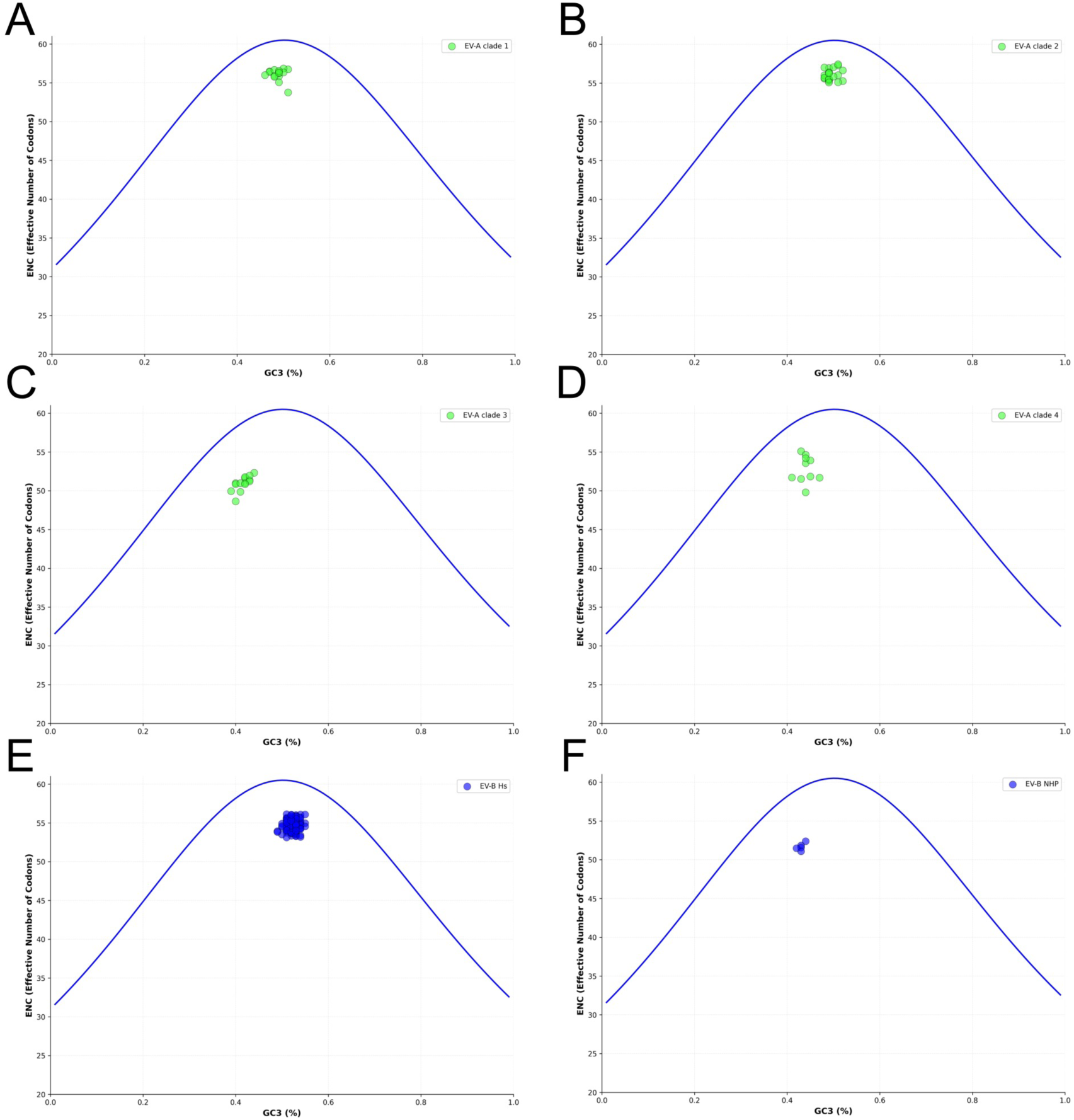
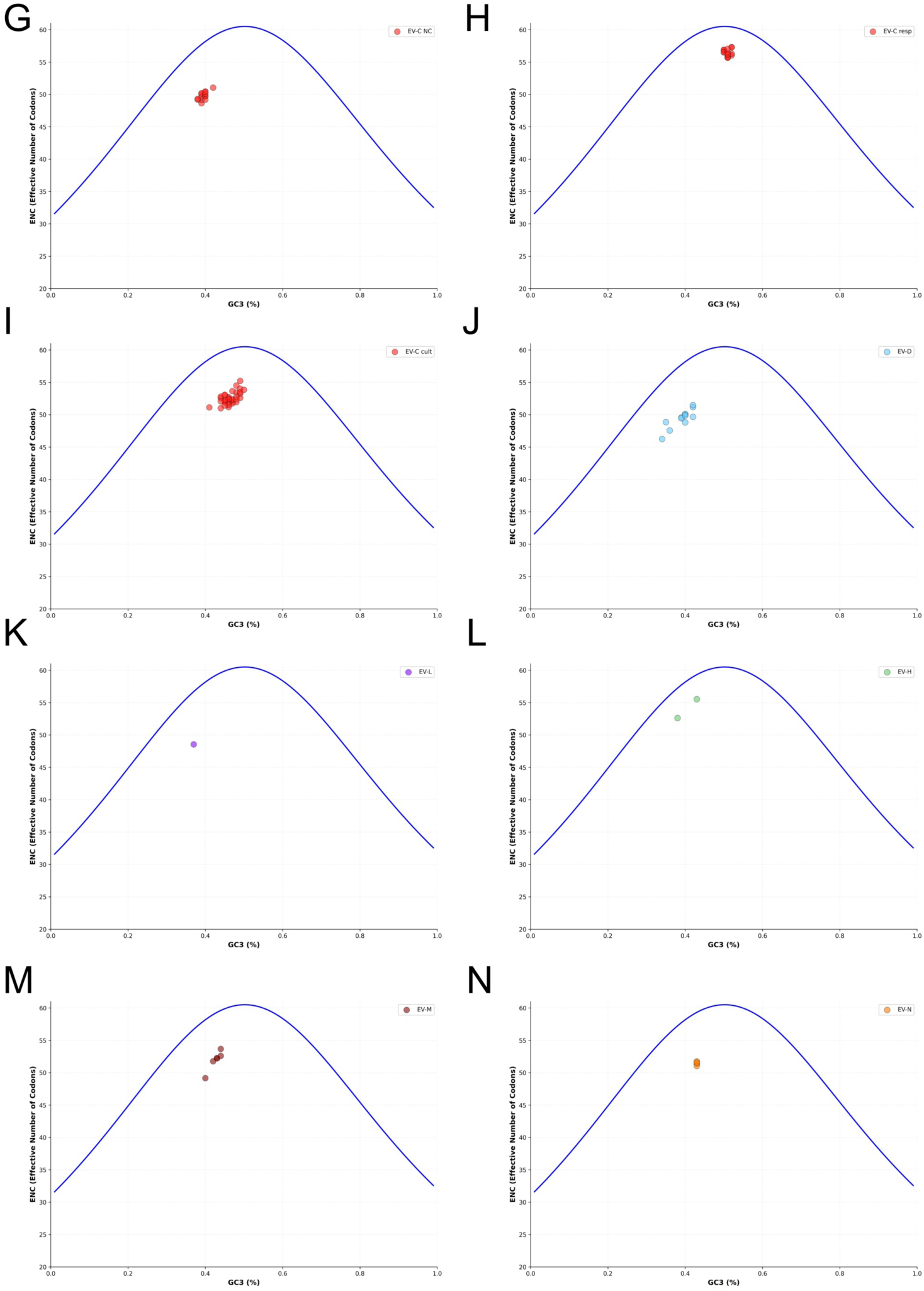
ENC–GC3 plots to determine bias mediated by GC3 position. The blue curve indicates the expected ENC values if GC3 composition alone drives codon usage bias. **(A)** EV-A clade 1 species. **(B)** EV-A clade 2 species. **(C)** EV-A clade 3 species. **(D)** EV-A clade 4 species. **(E)** Human EV-B. **(F)** NHP EV-B. The blue curve indicates the expected ENC values if GC3 composition alone drives codon usage bias. **(G)** Non-cultivable EV-C. **(H)** Respiratory EV-C. **(I)** Cultivable EV-C. **(J)** EV-D. **(K)** EV-L. **(L)** EV-H. **(M)** EV-M. **(N)** EV-N. Resp, Respiratory; NC, Non-cultivable; Cult: cultivable.

